# CB2 cannabinoid receptor-specific therapeutic antibody agonists for treatment of chemotherapy-induced peripheral neuropathy

**DOI:** 10.1101/2025.11.26.690750

**Authors:** Carlos Henrique Alves Jesus, Raghavender Gopireddy, Emily Sizemore, Jonah L. Wirt, Swastik Sen, Richard Yu, Toshihiko Takeuchi, Lauren Schwimmer, Andrea G. Hohmann

**Affiliations:** Department of Psychological and Brain Sciences, Indiana University, Bloomington, IN, USA; Gill Institute for Neuroscience, Indiana University, Bloomington, IN, USA; Abalone Bio Inc., Emeryville, CA; Program in Neuroscience, Indiana University, Bloomington, IN, USA

**Keywords:** endocannabinoid, neuropathic pain, paclitaxel, breast cancer, THC

## Abstract

Chemotherapy-induced peripheral neuropathy (CIPN) is a debilitating complication of cancer treatment. CB2 cannabinoid receptor activation reduces inflammation and is an attractive therapeutic target. Antibodies targeting G protein-coupled receptors (GPCRs), like CB2, offer high specificity and peripheral-restriction, thereby minimizing off-target activity. Here, we investigated the efficacy of first-in-class CB2-specific antibody agonists (AB110 and AB120) and an isotype control (AB100) on mechanical and cold hypersensitivity induced by paclitaxel in both tumor-free and mammary (4T1) tumor-bearing female mice. These CB2 antibody agonists exhibit biased G-α signaling and also reduce macrophage markers and pro-inflammatory cytokines *in vitro*. Paclitaxel produced behavioral hypersensitivities to mechanical and cold stimulation, which were reduced by AB110 and AB120 for approximately 48 hours post-injection in female mice. Repeated daily dosing did not lead to tolerance to the anti-allodynic effects. Prophylactic treatment with AB110 and AB120 during paclitaxel treatment delayed, but did not prevent, the development of paclitaxel-induced behavioral hypersensitivities after termination of dosing with antibody agonists. AB100 had no effect under any conditions. The anti-allodynic effects of AB120 were absent in CB2 knockout mice, confirming pharmacological specificity via CB2 receptors. Furthermore, AB120 remained effective in paclitaxel-treated tumor-bearing mice. Neither AB110 nor AB120 affected locomotor activity in otherwise naïve mice. The cytotoxic activity of paclitaxel on 4T1 tumor cell line was maintained in the presence of CB2 antibody agonists *in vitro*. Overall, our results suggest that CB2-specific antibody agonists are promising candidates for treating CIPN, providing lasting pain relief without tolerance, off target effects or unwanted CB1-mediated motor side effects.

## 1. Introduction

Chemotherapy remains an effective anti-cancer treatment but is often accompanied by complications [1], including chemotherapy-induced peripheral neuropathy (CIPN). CIPN is associated with known chemotherapeutic agents, including platinum-based compounds, taxanes, and vinca alkaloids [59]. Over 80% of patients undergoing chemotherapy develop CIPN, marked by numbness, tingling, and/or pain, all of which diminish quality of life [5]. Duloxetine remains the only pharmacological agent with demonstrated efficacy, albeit modest, for managing CIPN, and other medications (e.g., anticonvulsants, antidepressants, opioids) are frequently used off-label [17; 59]. Tolerance, dependence liability, sedation, and adverse side-effects limit utility of current available therapies [43]. Therefore, identification of safer, more effective treatments is of critical importance.

The endocannabinoid system contains multiple targets for analgesic drug development [55; 60]. However, direct activation of CB1 cannabinoid receptors in the central nervous system (CNS) produces unwanted side effects [55; 60]. By contrast, activation of CB2 cannabinoid receptors reduces pain behaviors in preclinical pain models [22; 24; 37-39; 47], without producing adverse CNS side effects. This is primarily due to the limited expression of CB2 receptors, which are predominantly found in immune cells, in the CNS [3; 51]. Our group has previously shown that therapeutic efficacy of small molecule CB2 agonists in neuropathic and inflammatory pain models depends on the expression of these receptors in peripheral sensory neurons [7; 22; 24]; efficacy was absent in Advillin^CRE/+^; CB ^f/f^ conditional knockout (cKO) mice [7; 22; 24]. Furthermore, treatment with paclitaxel dynamically increased CB2 expression, as defined by EGFP reporter, in Langerhans cells and in keratinocytes in paw skin of mice [39], supporting peripheral mechanisms in the analgesic effects of CB2 receptor activation.

Despite promising preclinical findings, CB2 agonists present limitations, including off-target effects [66], rapid clearance [71] and potential unintended activation of CB1 receptors via functional receptor heteromers [6]. Clinical trials on CB2 agonists not typically targeted therapeutic conditions supported by preclinical literature and have yielded limited results [9; 50]. These challenges underscore the need for novel strategies to more effectively target CB2 receptors and improve therapeutic outcomes in pain management. Therapeutic antibodies can precisely target molecules involved in diseases such as cancer, autoimmune disorders and infectious diseases [42], offering potential advantages over small molecule agonists through longer-lasting, peripheral, and highly specific effects [67]. However, developing therapeutic antibodies targeting G-protein coupled receptors (GPCRs) remains a challenge due to their complex structure and dynamic conformation [32].

Here, we developed first-in-class CB2-specific therapeutic antibody agonists, AB110 and AB120, and evaluated them for therapeutic efficacy in a preclinical model of paclitaxel-induced CIPN. These antibodies are CB2-specific and Gα-biased, and suppress the expression of macrophage activation markers and key pro-inflammatory cytokines, including IL-6, IL-1β, and TNF-α [21]. Specifically, we examined whether these antibodies could reduce paclitaxel-induced mechanical and cold hypersensitivities in comparison to an inactive isotype control, AB100. We assessed whether repeated dosing would induce tolerance and whether prophylactic treatment could prevent CIPN. Lastly, to enhance translational relevance, we asked whether our lead CB2-targeting antibody would suppress paclitaxel-induced CIPN in mice bearing mammary carcinoma.

## 2. Methods

### 2.1 Subjects

All experiments were conducted using 12-week-old female mice on a C57BL/6 background (Jackson Laboratories). Mammary carcinoma model was induced in 12-week-old female mice on a BALB/c background (Jackson Laboratories). Adult CB_2_KO female mice (B6.129P2-CNR2 (tm1Dgen/J)) on C57BL/6J background were bred at Indiana University under DA047858 (to AGH and Ken Mackie). Mice were housed under standard environmental conditions, including a temperature of 21 ± 2°C, 45% humidity, and a 12-hour light/dark cycle. All experimental procedures were approved by the Bloomington Institutional Animal Care and Use Committee at Indiana University (BIACUC) and were conducted in accordance with the animal care guidelines established by the International Association for the Study of Pain.

### 2.2 Drugs and Chemicals

Paclitaxel (Tecoland Corporation, Irvine, CA) was prepared in a vehicle composed of Cremophor (Sigma-Aldrich, St. Louis, MO), ethanol (Sigma-Aldrich), and saline (Aquilite System, Hospira Inc., Lake Forest, IL) mixed in a 1:1:18 ratio, following previously established methods [70]. CP55,940 was purchased from Cayman Pharmaceuticals and prepared in a vehicle composed of dimethyl sulfoxide (DMSO, Sigma-Aldrich), ethanol, emulphor (Sigma-Aldrich, St. Louis, MO) and saline in a ratio of 5:2:2:16 [14]. NKH 477, HU-308 and RO-1276 were purchased from Tocris. 3-Isobutyl-1-methylxanthine (IBMX), LPS and IL-4 were purchased from Sigma-Aldrich. SR144528 was purchased from Tocris Bioscience.

### 2.3 Antibody cloning and production

Antibody cloning and production was performed as previously described [21]. Briefly, VHH nucleotide sequences for AB100, AB110, and AB120 were synthesized as gene fragments by IDT and cloned into the TGEX-Fc expression vector (Antibody Design Labs) using NEBuilder HiFi DNA Assembly (New England Biolabs). Each construct was transfected into Expi293F cells (Thermo Fisher Scientific) using the Expi293 transfection reagent, following the manufacturer’s protocol. Cells were incubated at 37 °C with 8% CO_₂_ for 5 days. VHH-Fc fusion proteins were purified from the culture supernatant using MabSelect Sure protein A resin (Cytiva) via gravity flow. Purified antibodies were dialyzed into 1X PBS (pH 7.4). AB100, AB110 and AB120 were freshly diluted in phosphate-buffered saline (PBS, pH 7.4) before intraperitoneal (i.p.) injections in mice.

### 2.4 Cell culture for in vitro signaling assay

Mouse macrophage cell line (RAW264.7) was purchased from the American Type Culture Collection and cultured in Dulbecco’s modified Eagle medium (DMEM) from ATCC supplemented with 10% FBS, 20mM HEPES and 1X antibiotic and antimycotic at 37°C in a humidified incubator with 5% CO_2._Cells were maintained in a humidified incubator and routinely passaged when they reach approximately 75-95% confluency to prevent overgrowth. Prior to passaging, cells were washed with phosphate-buffered saline (PBS) and detached carefully using cell scrapers (Avantor). After detachment, cells were resuspended in fresh medium and either replated at the desired density or collected for downstream application.

### 2.5 Luminescence cAMP assay

cAMP-Glo Max assay kit (Promega, V1682) was used to measure cAMP signals in RAW264.7 cells as described previously [21]. Briefly, 384 well view plate (Revvity) was coated with Poly-D-lysine (Sigma Aldrich) and washed with twice with PBS. RAW264.7 cells were seeded (10,000 cells per well) onto the plate and incubated 16-18 hours at 37°C with 5% CO_2_. Cells were treated with test compounds, briefly centrifuged, and incubated for 30 minutes at 37°C. For experiments evaluating mediation by CB2, CB2 agonists were applied (EC80, 1uM) concurrently with CB2 antagonist/inverse agonist SR144528. After 30 minutes, PKA cAMP detection reagent (Glo one) with PKA was added to cells, briefly centrifuged, and incubated 20 minutes at ambient temperature (RT). In the final step, kinase Glo reagent was added to the cells, briefly centrifuged, and incubated for 10 minutes at RT. A Synergy HTX multi-mode microplate reader (Bio Tek) was used to detect the luminescence signal. The data were fit with a non-linear regression of variable slope using GraphPad Prism software (version 10.0).

### 2.6 β*-arrestin-2 recruitment assay*

The β-arrestin-2 recruitment assay was described previously [21]. Briefly, HTRF β-arrestin 2 Recruitment Detection Kit (Revvity) was used to measure β-arrestin-2 recruitment. RAW264.7 cells were seeded on 96 well culture plate and incubated for 24 hours at 37 °C with 5% CO_2_. After 24 hours, cells were transiently transfected with β-arrestin (20ng) and empty pcDNA3 (150ng) plasmids with Lipofectamine 2000 transfection kit (Thermo Fisher Scientific) and incubated for 24 hours at 37°C and 5% CO_2_. The media was removed after 24 hours and cells were treated with test compounds and incubated for 30 minutes at RT. Later, the media was removed, and stabilization buffer was added to the plate and incubated for 15 minutes at RT. Stabilization buffer was removed, and the plate was washed 3x with wash buffer. Afterwards, pre-mixed d2 and Eu cryptate antibodies were added to the cells and incubated overnight at RT. Next day, a Victor2 multimode plate reader was used read HTRF signal. The ratio ((em.655/em615)*10,000) was calculated and the data fit with a non-linear regression of variable slope using GraphPad Prism software (version 10.0).

### 2.7 Quantitative PCR

Macrophage gene expression by q-PCR was described previously [19; 25; 40]. Briefly, cells were seeded on 6-well culture plates and incubate 24 hours at 37°C with 5% CO_2_. Next, cells were serum starved for 24 hrs, test compounds were added to the cells and incubated 16 to 18 hours at 37°C with 5% CO2. After 18 hours, cells were incubated with LPS (100ng/ml) or IL-4 (10ng/ml) for 8 hours and 16 hours, respectively. After incubation, cells were washed in PBS and lysed. Total RNA was extracted from cells using RNeasy Mini Kit (Qiagen) according to the manufacturer’s protocol. Afterwards, cDNA was synthesized using Super script IV VILO master mix kit (Invitrogen). Finally, quantitative PCR (qPCR) was performed on AriaMx RT-PCR system (Agilent), using Power SYBR Green Universal Master mix (Applied Biosystems). The relative expression level of specific mRNA was determined by the comparative cycle threshold (CT) method (2^-ΔΔCT^), normalized to the endogenous control gene GAPDH. Each RNA sample was assayed in triplicate. The primers used in real-time PCR are listed in Supplemental Table 1.

### 2.8 Cell culture

The 4T1 murine mammary carcinoma cell line was kindly provided by Dr. Harikrishna Nakshatri from Indiana University School of Medicine in Indianapolis, Indiana. The HEK293 cell line was kindly provided by Dr. Ken Mackie, Gill Institute for Neuroscience, Indiana University Bloomington. The cell lines were cultured according to previous studies [34; 74], with minor modifications. Briefly, 4T1 cells were cultured in RPMI-1640 medium (Thermo Fisher Scientific, United States, A1049101) supplemented with 10% fetal bovine serum (Fisher Scientific, United States, A5670701) and 1% penicillin-streptomycin (Thermo Fisher Scientific, United States, 15140122) under standard cell culture conditions (37°C, 5% CO_₂_). Cells were maintained in a humidified incubator and routinely passaged when they reach approximately 75-95% confluency to prevent overgrowth. Prior to passaging, cells were washed with phosphate-buffered saline (PBS) and detached using 0.05% trypsin-EDTA (Fisher Scientific, 25-300-062). After detachment, cells were resuspended in fresh medium and either replated at the desired density or collected for downstream application.

### 2.9 MTT cell viability assay and synergy reference models

4T1 and HEK293 cells (see section 2.9 for cell cuture methods) were incubated at 37 °C in a humidified atmosphere containing 5% CO_2_. Cells were plated at a density of 3000 cells per well in standard flat-bottomed 96-well plates. Cell viability was assessed using the MTT assay as previously described [36; 65; 70]. Briefly, each CB2-specific antibody candidate was tested in an 8 × 8 dose-response matrix, both alone and in combination with increasing concentrations of paclitaxel. Concentrations ranged from 0–50 µM for the CB2-specific antibodies and 0–500 nM for paclitaxel. After a 72-hour incubation period, 10 µL of MTT solution was added to each well. Formazan crystals were solubilized using 100 µL of SDS/HCl solubilization buffer, and absorbance was measured at 570 nm. Cell viability was calculated by comparing absorbance values of treated wells to untreated controls and expressed as a percentage of control. Data represent the mean of three (4T1) or four (HEK293) independent experiments. Dose–response curves were normalized and analyzed using nonlinear regression to determine EC_₅₀_ values. Drug interaction analysis was conducted using Combenefit opensource software (Cancer Research UK Cambridge Institute, Cambridge, UK) and SynergyFinder (https://synergyfinder.fimm.fi). These analyses enabled comparison of the effects of combining AB100, AB110, or AB120 with paclitaxel on the viability of 4T1 tumor cells and non-tumor HEK293 cells. We employed three synergy models: Highest Single Agent (HSA), Bliss independence and Loewe additivity. The Highest Single Agent (HSA) model assumes that the expected effect of a drug combination is equal to the greater of the individual effects observed at the same concentrations [28]. The Bliss independence model is based on the premise that the two drugs act independently, and therefore, the predicted combination effect is derived from the probability of their separate actions occurring together [28]. The Loewe additivity model estimates the expected outcome under the assumption that both drugs behave as if they are identical in their mechanism of action [28].

### 2.10 Paclitaxel-induced neuropathic nociception

Paclitaxel (4 mg/kg/day) or its Cremophor-based vehicle was administered (i.p.) once daily on alternating days, for a total of four injections. Mechanical and cold responsiveness were evaluated at baseline (BL, prior to the first injection) and subsequently every 3 to 5 days following the start of paclitaxel treatment, continuing for up to 14 days prior to antibody treatments.

### 2.11 Mammary carcinoma cell inoculation and paclitaxel regimen in tumor bearing mice

Mammary carcinoma tumors were established in female BALB/c mice following a previously described method [74], with minor modifications. In brief, to generate tumor-bearing mice, after 3 passage protocols, viable 4T1 mammary carcinoma cells were suspended in a sterile 1:1 mixture of PBS and Matrigel basement membrane matrix (Corning, Glendale, Arizona, USA; 354248). A concentration of 1×10 cells in 100 µL was orthotopically injected unilaterally into the second mammary fat pad of syngeneic, immunocompetent female BALB/c mice. This immunocompetent model allowed tumor development in a physiologically relevant immune environment. All injections were carried out under sterile conditions and light anesthesia (5% isoflurane) to reduce animal stress. Tumors became palpable within 4–5 days post-inoculation and were routinely monitored using calipers. Tumor volume was calculated using the formula: [(width² × length) / 2]. Paclitaxel (10 mg/kg, i.p.) was given once daily, every other day for a total of three injections. Mechanical and cold sensitivity responses were evaluated at baseline (BL, prior to the first injection), at the day where tumors became palpable (day 0, D0) and subsequently on days 2, 4, 5 (antibody testing day) and 6 (24-hour post antibody treatment).

### 2.12 Assessment of mechanical paw withdrawal thresholds

Paw withdrawal thresholds were assessed using methods previously established by our research group [7; 14-16; 22-24; 38; 39; 70]. In brief, animals were acclimated for 30 minutes in individual inverted plexiglass chambers placed on a metal mesh platform supported by a stable wooden table. Paw withdrawal thresholds were measured using a semi-flexible 90-gram probe attached to an electronic von Frey analgesiometer (IITC Life Science, Woodland Hills, CA). The mechanical withdrawal threshold was defined as the unit (in grams) required to elicit a paw withdrawal response. Measurements were taken in duplicate for each hind paw, with a 2-minute interval between tests to prevent sensitization. The values from both paws were then averaged to yield a single threshold per animal per time point, based on previous findings indicating that the paclitaxel dosing protocol induces symmetrical neuropathy with no significant difference between paws.

### 2.13 Assessment of cold responsiveness to acetone

Cold sensitivity was assessed immediately following the evaluation of mechanical paw withdrawal thresholds in the same mice, as previously described [37–39; 70]. A 1 mL syringe (with the needle removed) was filled with acetone (Sigma-Aldrich). To apply the stimulus, a drop of acetone was gently formed at the syringe tip by pressing the plunger and carefully placed onto the plantar surface of the hind paw, ensuring no direct contact with the syringe to avoid mechanical stimulation. The duration (in seconds) that the mouse attended to the stimulated paw, through behaviors such as lifting, biting, licking, shaking, or flinching, was recorded three times per paw, with several minutes between applications to prevent sensitization. The response durations were then averaged across both paws for each animal.

### 2.14 Assessment of motor ataxia and locomotor activity

For the rotarod assay, mice first underwent two consecutive days of training. During each training day, mice were required to remain on the rotarod apparatus (IITC Life Sciences Inc.) for a minimum of 30 seconds across three consecutive trials. On the third day, baseline performance was recorded before animals were randomly assigned to drug treatment. Post-treatment assessments were conducted 2 hours after injection, with latency to descend recorded both at baseline (BL) and following pharmacological manipulations. For locomotor activity assessments, the same mice from the rotarod experiment underwent a 2-week washout period and then were placed in activity monitoring arenas (16 × 16 × 12 inches; Omnitech Superflex Nodes, Omnitech, Columbus, OH), with all movements automatically tracked using Fusion 6.5 software. The testing environment was maintained at ∼80 lux using tungsten lighting, and white noise (62–63 dB) was continuously played to minimize external disturbances. On day 0, mice were habituated to the arena for 15 minutes. Baseline locomotor activity was recorded on day 1 without injection. On day 2, mice received the same treatments administered in the rota-rod test and locomotor activity was assessed 2 hours post-injection. Total distance traveled as well as rest time were analyzed for both baseline and post-injection test sessions that were each 30 minutes in duration [70].

### 2.15 Experimental procedures

Our previous studies using small-molecule CB2 agonists did not reveal any sex-dependent differences in potency or efficacy in neuropathy models induced by paclitaxel or the antiretroviral agent dideoxycytidine [7; 24; 39]. Therefore, female mice were used exclusively to evaluate CB2 antibody agonists, reflecting the therapeutic focus on treating CIPN in female breast-cancer survivors [64]. The therapeutic effectiveness of specific CB2 antibody agonists was assessed in a paclitaxel-induced peripheral neuropathy model in female mice, using the following treatment schedules and experimental design:

#### Experiment 1: Development of paclitaxel-induced mechanical and cold hypersensitivity

We assessed the development of mechanical and cold hypersensitivity in female mice following paclitaxel treatment, administered intraperitoneally at a dose of 4 mg/kg (i.p.) every other day for a total of four injections. Mechanical thresholds and response to acetone applied to the hind paw were measured at baseline and on days 4, 7, 11, and 14 after the initial paclitaxel dose.

#### Experiment 2: Effect of CB2-specific antibody agonist AB110 on paclitaxel-induced mechanical and cold hypersensitivity in female mice

We evaluated antinociceptive effects of different doses of the CB2 antibody agonist AB110 (5 or 25 mg/kg, i.p.) and compared it to the inactive isotype control AB100 (25 mg/kg, i.p.) or vehicle PBS (i.p.). All treatments were administered 14 days after the initial paclitaxel treatment, when mechanical and cold hypersensitivity were fully developed and stable. Mechanical thresholds and response to acetone applied to the hind paw were measured at 0.5, 2, 4, 24, 48, and 72 hours following antibody administration.

#### Experiment 3: Effect of CB2-specific antibody agonist AB120 on paclitaxel-induced mechanical and cold hypersensitivity in female mice

We assessed the dose response antinociceptive effects of the CB2 antibody agonist AB120 (5 or 25 mg/kg, i.p.) and compared its efficacy to that of the antibody candidate AB110 (5 mg/kg, i.p.), or vehicle control PBS (i.p.) at 14 days following the initial paclitaxel administration. Mechanical thresholds and response to acetone were re-evaluated at 0.5, 2, 4, 24, 48, 72, and 96 hours after antibody treatment.

#### Experiment 4: Effect of repeated administration with CB2 antibody agonists AB120 and AB110 on paclitaxel-induced mechanical and cold hypersensitivity in female mice

We investigated whether repeated administration of the CB2 antibody agonist AB120 (5 or 25 mg/kg, i.p.) would lead to tolerance to analgesic efficacy. Separate groups received the CB2 agonist AB110 (5 mg/kg, i.p.), and vehicle PBS (i.p.). Female mice received daily injections of the antibody agonists for seven consecutive days (days 0–6). Mechanical thresholds and response to acetone were reassessed on days 1, 2, 5, and 7 following the initial antibody dose, always before the daily dosing.

#### Experiment 5: Effect of prophylactic administration with CB2 antibody agonist AB120 on the development of paclitaxel-induced mechanical and cold hypersensitivity in female mice

We examined whether the CB2 antibody agonist AB120 (5 or 25 mg/kg, i.p.), the isotype control AB100 (5 mg/kg, i.p.), or vehicle (PBS), administered once daily (i.e., 2 hours before each paclitaxel (4 mg/kg, i.p., given every other day for a total of four doses) injection and on intervening non-paclitaxel days), could prevent the development of mechanical and cold hypersensitivity in female mice. Mechanical thresholds and response to acetone were assessed at baseline (BL) and on days 2, 4, 6, and 8 following the initial antibody and paclitaxel treatment. Antibody administration was halted on day 8, and further assessments were conducted on days 11 and 15 to determine the durability of any therapeutic effects.

#### Experiment 6: Effect of CB2-specific antibody agonist AB120 in global CB2 knock out female mice

We investigated whether the antinociceptive efficacy of the CB2 antibody agonist AB120 is dependent on CB2 receptor expression by testing its effects in CB2 knockout (KO) mice. AB120 (5 mg/kg, i.p.) or vehicle (PBS) was administered to CB2 KO mice, while wild-type mice received AB120 (5 mg/kg, i.p.) as a control. Mechanical thresholds and response to acetone were assessed at baseline (BL), 14 days after the initial paclitaxel (PAX) regimen, and at 0.5, 2, 4, 24, 48, 72, and 96 hours following antibody administration.

#### Experiment 7: Effects of CB2-specific antibody agonist AB120 on paclitaxel-induced mechanical and cold hypersensitivity in mice with 4T1-induced mammary carcinoma

Female BALB/c mice were first assessed for baseline mechanical thresholds and response to acetone. Then they were then orthotopically inoculated with 4T1 cells into the second mammary fat pad. Once palpable mammary tumors developed, mechanical and cold sensitivity were re-evaluated to rule out tumor-induced hypersensitivity. Paclitaxel (10 mg/kg, i.p.) was subsequently administered every other day for a total of three injections on days 0, 2, and 4 following initial tumor detection. Sensory assessments were repeated prior to each paclitaxel dose on days 2 and 4, and again on day 5, before administration of the CB2 antibody agonist AB120 (5 mg/kg, i.p.) or the inactive isotype control AB100 (5 mg/kg, i.p.). Mechanical thresholds and acetone response were then measured at 0.5, 2, 4, and 24 hours post-antibody treatment. Tumor growth was monitored throughout the study via caliper measurements.

#### Experiment 8: Effect of CB2-specific antibody agonists AB110 and AB120 on motor ataxia and locomotor activity in female mice

Female mice were evaluated for motor coordination and ataxia using the rotarod test following treatment. After completing training, qualification, and baseline trials, mice were randomly assigned to one of the following i.p. treatment groups: PBS, AB100 (5 mg/kg), AB110 (5 mg/kg), AB120 (5 or 25 mg/kg), or the synthetic cannabinoid agonist CP55940 (0.3 mg/kg). Two hours post-treatment, mice were placed on the rotarod apparatus, and latency to descend was recorded across two trials. The average latency was used as an index of motor performance and potential ataxia. To assess locomotor activity, mice underwent a two-week washout period following the rotarod drug experiment. After habituation (day 0) and baseline assessment (day 1), female mice received i.p. injections of either PBS, AB100 (5 mg/kg), AB110 (5 mg/kg), AB120 (5 or 25 mg/kg), or the synthetic cannabinoid agonist CP55940 (0.3 mg/kg) on day 2. Two hours post-treatment, mice were placed in an open-field activity arena for 30 minutes. Total distance traveled and time spent at rest were recorded as indicators of potential locomotor impairment induced by the treatments.

### 2.11 Statistical analysis

Mechanical paw withdrawal thresholds, response to acetone application, rotarod descent latency, total distance traveled, and rest time in the activity meter were analyzed using Two-way repeated measures ANOVA. Bonferroni’s *post hoc* test was applied for multiple comparisons. Two-tailed paired t-tests were applied to evaluate differences in paw withdrawal threshold and response to acetone between baseline (BL) and first day of palpable tumors (D0) in the mammary tumor bearing mice experiment. Planned comparison one-tailed t-tests were used to compare CP55,940 to BL and vehicle treatment in the rota-rod test. All statistical analyses were performed using GraphPad Prism 10 (GraphPad Software, La Jolla, CA), with a significance level set at *p* < 0.05. 3.

## 3. Results

### 3.1 CB2 specific antibody agonists inhibit cAMP production and lead to β-arrestin-2 recruitment in vitro

We employed a bioluminescence-based high-throughput assay to quantify cAMP levels. In this assay, RAW264.7 cells were stimulated with NKH477 (a forskolin derivative) to induce cAMP production and subsequently treated with AB110, AB1120, HU-308 (a CB2 small-molecule agonist), or AB100 (a negative isotype control antibody). Notably, both AB110 and AB120, but not AB100, strongly inhibited cAMP production in this mouse macrophage cell line. The EC_50_ and E_max_ of both antibodies are comparable to HU-308, a selective CB2 receptor agonist (**Figure 1A**, **Supplemental Table 2**). The agonist activity of AB110 and AB120 was inhibited *in vitro* by SR144528, a selective CB2 antagonist/inverse agonist (**Figure 1B, Supplemental Table 2**). Moreover, a TR-FRET-based high-throughput assay was employed to evaluate β-arrestin-2 recruitment by detecting the interaction between β-arrestin-2 and AP2, which associates with β-arrestin-2 only upon activation of CB2 receptors. Our results showed that AB110 and AB120 exhibited potencies comparable to HU-308 in promoting β-arrestin-2 recruitment in mouse macrophages (**Figure 1C, Supplemental Table 2**). Both antibodies showed partial agonist properties, with lower E_max_ when compared to HU-308 (**Figure 1C, Supplemental Table 2**).

**Figure 1.**
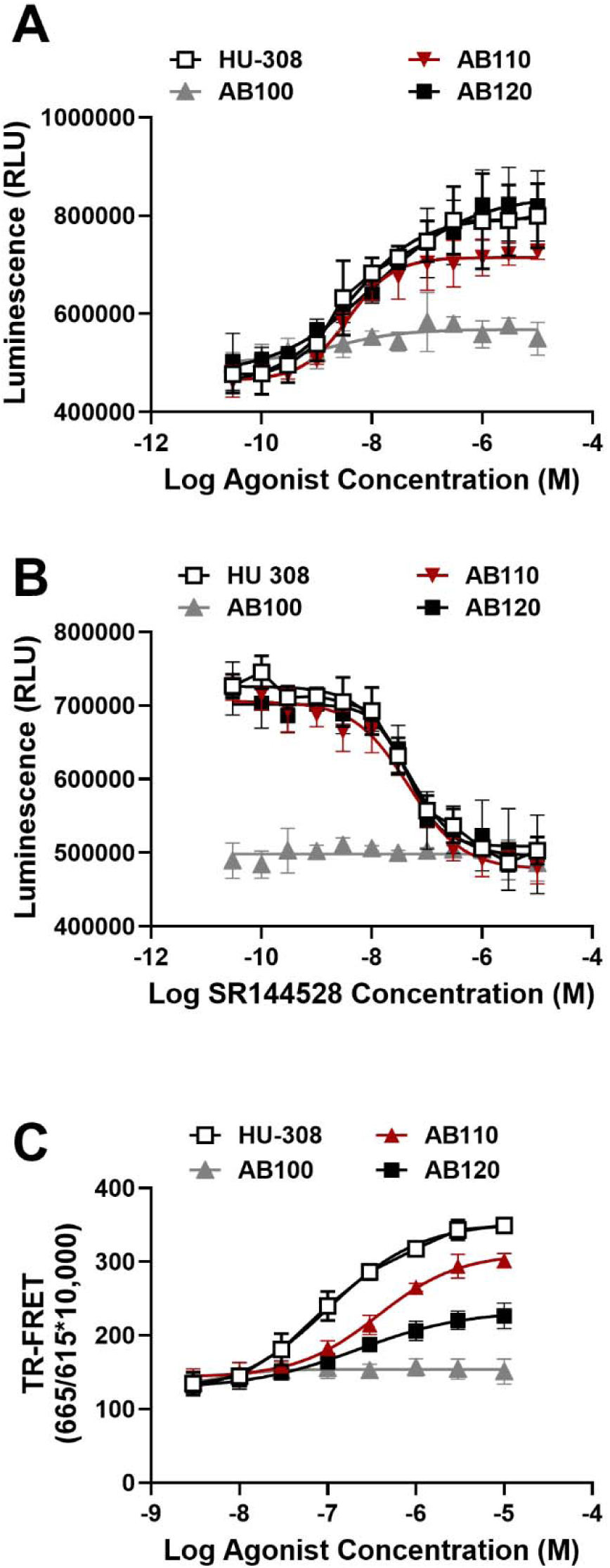
CB2 agonist antibodies inhibit cAMP production in RAW264.7 cells. The Bioluminescent cAMP-Glo™ Max Assay (Promega) was used to monitor cAMP levels where cAMP concentration is inversely proportional to luminescence signal. (**A**) CB2 specific agonist (HU-308) and CB2 specific agonist antibodies (AB110 and AB120) decrease NKH477-induced cAMP production. (**B**) The CB2-selective antagonist/inverse agonist SR144528 inhibits the agonist activity of HU-308, AB110, and AB120. (**C**) β-arrestin-2 recruitment by CB2 agonist antibodies in RAW264.7 cells. HTRF β-arrestin-2 recruitment assay was performed to measure β-arrestin-2 recruitment. RAW264.7 cells were transfected with 20 ng of human β-arrestin-2 expressing plasmid and 130ng of empty pcDNA3 plasmid and β-arrestin-2 recruitment was measured by TR-FRET. Data represents mean ± SD from four independent experiments.

### 3.2 CB2 agonist antibodies reduce mRNA expression associated with LPS and IL-4 activation of macrophages

LPS-stimulated RAW264.7 cells increased mRNA expression levels of pro-inflammatory markers in a manner that was sensitive to active CB2 agonists (HU308, AB110, AB120) but not the isotype control AB100 (**Figure 2A**, Treatment: F_5,60_ = 306.2, p<0.0001; Marker: F_4,60_ = 0.9331, p=0.4510; Interaction: F_20,60_ = 1.244, p=0.2526). LPS-stimulated cells showed consistent increased expression of pro-inflammatory markers, when compared to cells stimulated with buffer (p≤0.0011). CB2 antibody agonists, AB110 and AB120, decreased the expression of M1 macrophage associated pro-inflammatory mRNA (TNF-α, NOS2, IL-6, IL-1β and CCL4), to a level similar to that observed with the CB2 agonist HU-308, when compared with untreated LPS-stimulated cells (p<0.0001).

**Figure 2.**
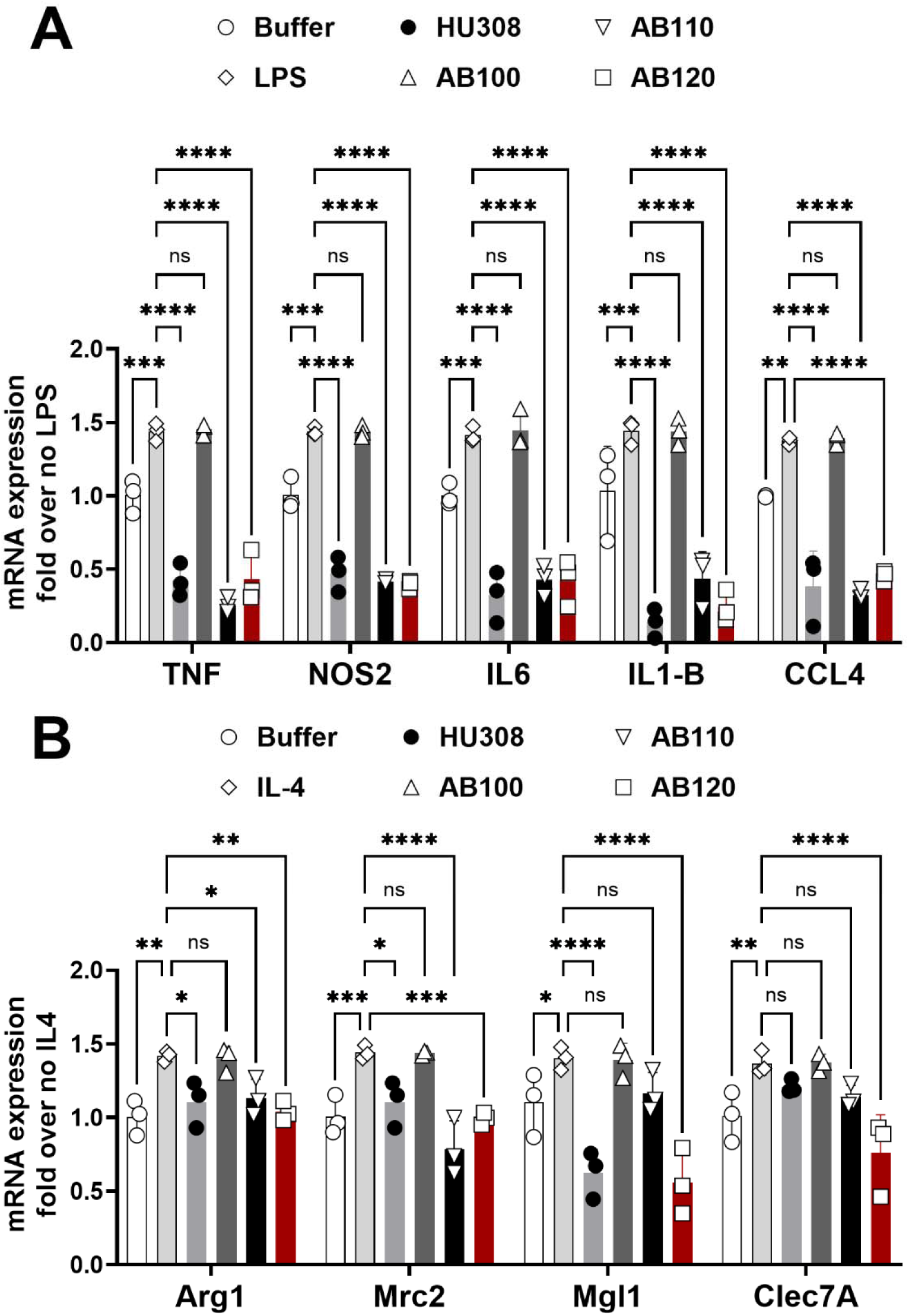
CB2 specific agonist antibodies reduce mRNA expression associated with LPS and IL-4 activation of macrophages. Serum starved RAW264.7 cells were treated with LPS (100ng/ml) for 8 hours or IL-4 (10ng/ml) for 16 hours. RT-PCR analysis for (**A**) M1 (LPS) associated gene expression and (**B**) M2 (IL-4) associated mRNA expression in RAW264.7 cells. Data represents mean ± SD from three independent experiments. Two-way ANOVA followed by Dunnet’s post hoc. Brackets: *\*P <0.05, **P <0.01, ***P <0.001 and ****P <0.0001*.

When RAW264.7 cells were treated with IL-4 to promote M2 polarization, mRNA expression levels of select markers were upregulated in a manner that was prevented by active CB2 agonists but not the isotype control AB100 (**Figure 2B**, Treatment: F_5,48_ = 35.98, p<0.0001; Marker: F_3,48_ = 3.665, p=0.0186; Interaction: F_15,48_ = 4.533, p<0.0001).

The expression of M2 macrophage-associated mRNA (Arg1, Mrc2, Mgl1, and Clec7A) increased (**Figure 2B**) after IL-4 stimulation, when compared to buffer control (p≤0.0335). CB2 agonist HU-308, and antibody agonists AB110 and AB120, inhibited the upregulation of Arg1 and Mrc2 in RAW264.7 cells (p≤0.0465). Only AB120 and HU-308 attenuated Mgl1 (p<0.0001). Only AB120 attenuated Clec7A expression in RAW264.7 cells (p<0.0001), when compared to control treated with IL-4 (**Figure 2B**). The antibody isotype control AB100 did not affect mRNA expression of any M1 or M2 markers when compared to LPS control (p≥0.99).

### 3.3 Paclitaxel produced hypersensitivity to mechanical and cold stimulation in mice

Paclitaxel treatment (4 mg/kg, i.p. administered on day 0, 2, 4 and 6) reduced paw withdrawal thresholds overall and in a time-dependent manner (**Figure 3A**; Treatment: F_1,7_ = 195.4, p<0.0001; Time: F_4,28_ = 14.09, p<0.0001; Interaction: F_4,28_ = 8.601, p=0.0001). Mechanical hypersensitivity was evident in paclitaxel-treated mice from days 4 to 14 (p<0.0001) following initiation of paclitaxel dosing compared to the Cremophor-based vehicle group. Similarly, responses to acetone application increased following paclitaxel treatment overall and in a time-dependent manner (**Figure 3B**; Treatment: F_1,7_= 185.8, p<0.0001; Time: F_4,28_= 22.82, p<0.0001; Interaction: F_4,28_ = 13.60, p<0.0001). Cold hypersensitivity was observed from day 4 through day 14 post-treatment (day 4: p=0.0002; days 7–14: p<0.0001).

**Figure 3.**
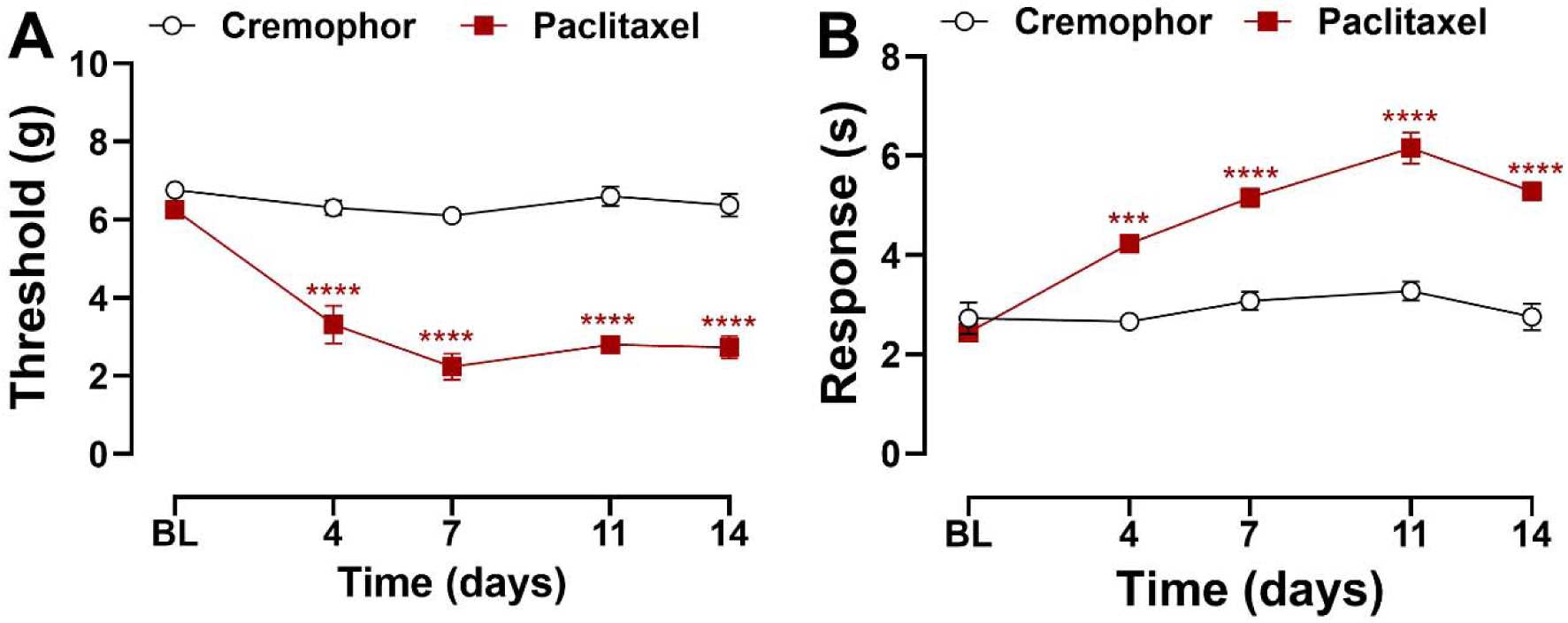
Paclitaxel induces mechanical and cold hypersensitivity in female mice. Mice treated with paclitaxel displayed (**A**) mechanical and (**B**) cold hypersensitivity 4 – 14 days following paclitaxel dosing regimen prior to antibody treatment. Data show mean (± SEM) (n=3-6). Two-way ANOVA followed by Bonferroni’s *post hoc* test. Interaction between time points and treatment effect: *\*\*\*P <0.001, ****P <0.0001* Cremophor vehicle vs. Paclitaxel.

### 3.4 CB2-specific antibody agonist AB110 attenuated paclitaxel-induced mechanical and cold hypersensitivity in mice

In paclitaxel-treated mice, AB110 increased paw withdrawal thresholds overall, withdrawal thresholds changed across time but an interaction between treatment and time was not observed (**Figure 4A**; Treatment: F_3,17_=67.55, p<0.0001; Time: F_3.969,67.48_=7.024, p<0.0001; Interaction: F_11.91,67.48_=1.665, p=0.0954). All paclitaxel-treated groups exhibited lower paw withdrawal thresholds compared to the Cremophor vehicle group (p<0.0001). AB110 at doses of either 5 mg/kg or 25 mg/kg (i.p.) attenuated paclitaxel-induced mechanical hypersensitivity compared to the PBS-treated control group (p<0.0001 for both doses). No difference was observed between the two AB110 doses (p>0.99).

**Figure 4.**
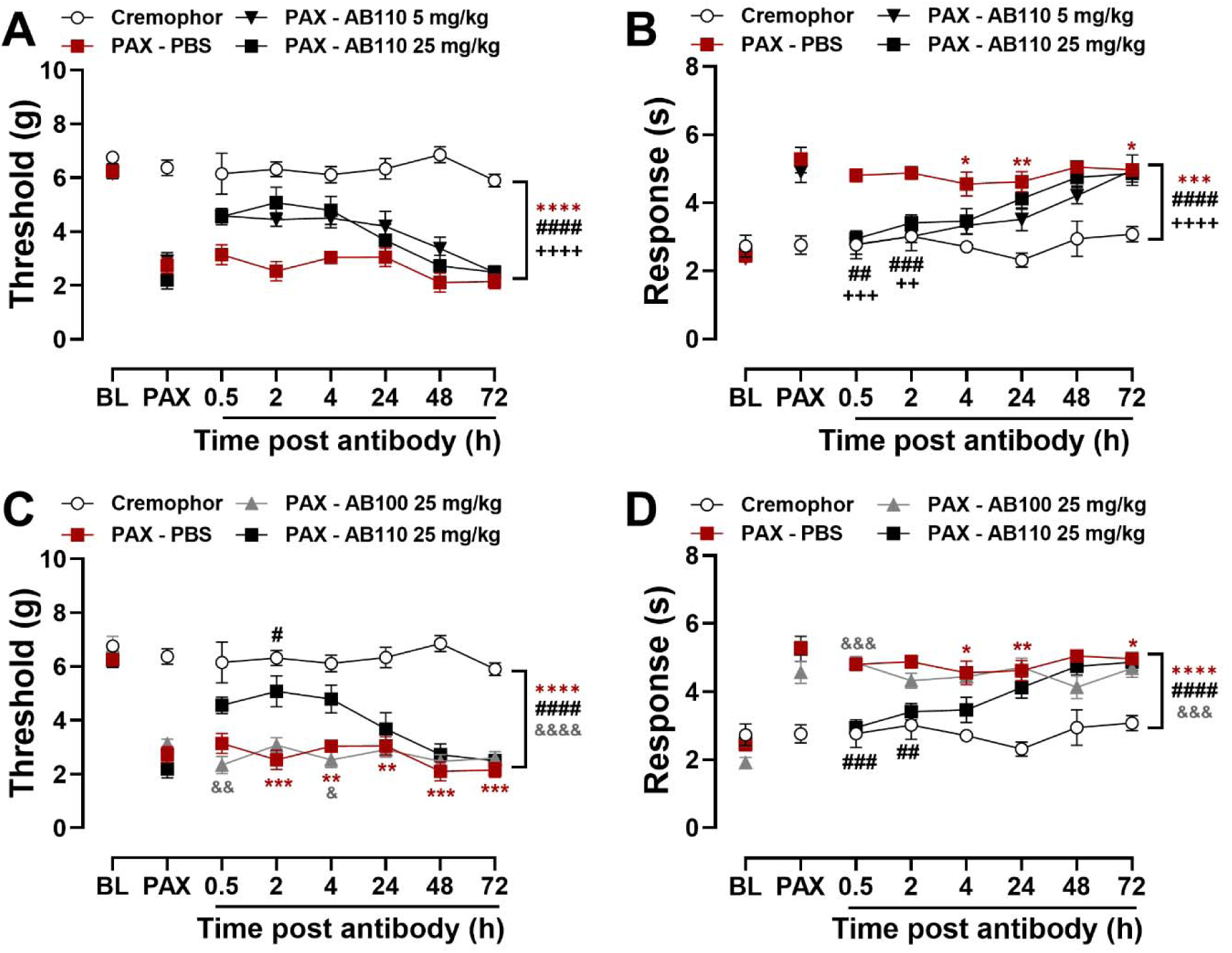
CB2-specific antibody agonist AB110 attenuates paclitaxel-induced mechanical and cold hypersensitivity in female mice. Mice were given AB110 or inactive isotype control AB100 at 14 days post initial paclitaxel dose, when CIPN was established. AB110 (5 and 25 mg/kg, i.p.) attenuated paclitaxel-induced (**A**) mechanical and (**B**) cold hypersensitivity. In comparison to inactive antibody AB100 (25 mg/kg, i.p.), AB110 (25 mg/kg, i.p.) attenuated (**C**) mechanical and (**D**) cold hypersensitivity induced by paclitaxel. Data show mean (± SEM) (n=3-6). Two-way ANOVA followed by Bonferroni’s *post hoc* test. Interaction between time and treatment: *\*P <0.05 **P <0.01, ***P <0.001 Cremophor vs. PAX-PBS; ^#^P <0.05, ^##^P <0.01, ^###^P <0.001 PAX-PBS vs. PAX-AB110 5 mg/kg; ^++^P <0.01, ^+++^P <0.001 PAX-PBS vs. AB110 25 mg/kg. ^&^P <0.05, ^&&^P <0.01 PAX-AB100 25 mg/kg vs. PAX-AB110 25 mg/kg. Brackets show main treatment effect: ***P <0.001; ****P <0.0001 Cremophor vs. PAX-treated groups; Panels A-B: ^####^P <0.0001 PAX-PBS vs. PAX-AB110 5 mg/kg; ^++++^P<0.0001 PAX-PBS vs. PAX-AB110 25 mg/kg; Panel C-D: ^####^ P<0.0001 PAX-PBS vs. PAX-AB110 25 mg/kg; ^&&&^P<0.001, ^&&&&^P <0.0001 PAX-AB100 vs. PAX-AB110*.

AB110 attenuated paclitaxel-induced cold hypersensitivity overall and in a time-dependent manner (**Figure 4B**: Treatment: F_3,17_=18.64, p<0.0001; Time: F_3.768,64.05_=12.03, p<0.0001; Interaction: F_11.30,64.05_=2.970, p=0.0028). Overall, paclitaxel increased responsiveness to acetone in comparison to the Cremophor vehicle-treated group (p<0.0001). AB110 suppressed paclitaxel-induced cold hypersensitivity (p<0.0001), with no difference observed between doses (p=0.38). Acetone-evoked responses were higher in PBS-treated, paclitaxel-exposed mice from 4-72 h (p≤0.0185) post-treatment compared to Cremophor vehicle-treated controls. AB110 (5 mg/kg, i.p.) reduced paclitaxel-induced cold hypersensitivity at 0.5 (p=0.0017) and 2 hours (p=0.0002) after injection. Similarly, AB110 (25 mg/kg, i.p.) attenuated paclitaxel-induced cold hypersensitivity at 0.5 (p=0.0002) and 2 (p=0.0045) hours post injection.

AB110 also suppressed paclitaxel-induced mechanical hypersensitivity relative to the inactive isotype control AB100 overall and in a time-dependent manner (**Figure 4C**: Treatment: F_3,17_=101.6, p<0.0001; Time: F_4.121,70.05_=3.480, p=0.0112; Interaction: F_12.36,70.05_=2.306, p=0.0145). Throughout the study, paclitaxel reduced mechanical thresholds compared to the Cremophor vehicle-treated control group (p<0.0001). AB110 consistently reduced mechanical hypersensitivity (p<0.0001), whereas AB100 showed no effect (p>0.99). Paclitaxel-treated mice receiving PBS exhibited lower paw withdrawal thresholds from 2-72 h (p≤0.0098) post-injection compared to Cremophor vehicle-treated controls. AB110 (25 mg/kg, i.p.) reliably attenuated paclitaxel-induced mechanical hypersensitivity at 2 hours post-treatment (p=0.0314), and this effect was also greater than that of the isotype control at 4 hours post-injection (p=0.0341).

Antibody treatment with AB110, but not AB100, reduced acetone responsiveness in a treatment and time-dependent manner (**Figure 4D**: Treatment: F_3,17_=22.53, p<0.0001; Time: F_3.658,62.19_=3.850, p=0.0090; Interaction: F_10.97,62.19_=3.731, p=0.0004). Paclitaxel-treated mice receiving PBS showed consistently elevated responses to acetone compared to Cremophor vehicle-treated controls across the observation interval (p<0.0001). AB110 attenuated paclitaxel-induced cold responsiveness overall (p<0.0001), whereas AB100 did not differ from treatment with PBS (p=0.29). Paclitaxel-treated mice receiving PBS exhibited increased acetone-evoked responses at 4-72 h post-injection (p≤0.0185) compared to Cremophor vehicle-treated controls. AB110 (25 mg/kg, i.p.) reliably reduced cold hypersensitivity from 0.5-2 h (p≤0.0045) post-injection. Cold responsiveness was lower in the AB110-treated group compared to the inactive isotype control AB100 at 0.5 hours post-treatment (p=0.0002).

### 3.5 CB2-specific antibody agonist AB120 attenuates paclitaxel-induced allodynia with superior efficacy compared to AB110

AB120 (5 and 25 mg/kg, i.p.) attenuated paclitaxel-induced mechanical hypersensitivity in a treatment and time-dependent manner (**Figure 5A**: Treatment: F_3,18_=32.14, p<0.0001; Time: F_4.436,79.84_=4.785, p=0.0011; Interaction: F_13.31,79.84_=2.074, p=0.0239). Paclitaxel-treated mice exhibited lower paw withdrawal thresholds than Cremophor vehicle-treated controls across the observation interval (p<0.0001). AB120 attenuated paclitaxel-induced mechanical hypersensitivity at all tested doses compared to PBS-treated mice (p<0.0001 for each dose). Paclitaxel-treated mice receiving vehicle (PBS) exhibited reduced paw withdrawal thresholds compared to the Cremophor control group at multiple time points (0.5-96 h: p≤0.0499). AB120 (5 mg/kg, i.p.) alleviated paclitaxel-induced mechanical hypersensitivity (2-48h: p≤0.0143) relative to PBS treatment. AB120 (25 mg/kg, i.p.) also showed efficacy at the 48-hour post-injection time point (p=0.0475), relative to PBS-treated mice.

**Figure 5.**
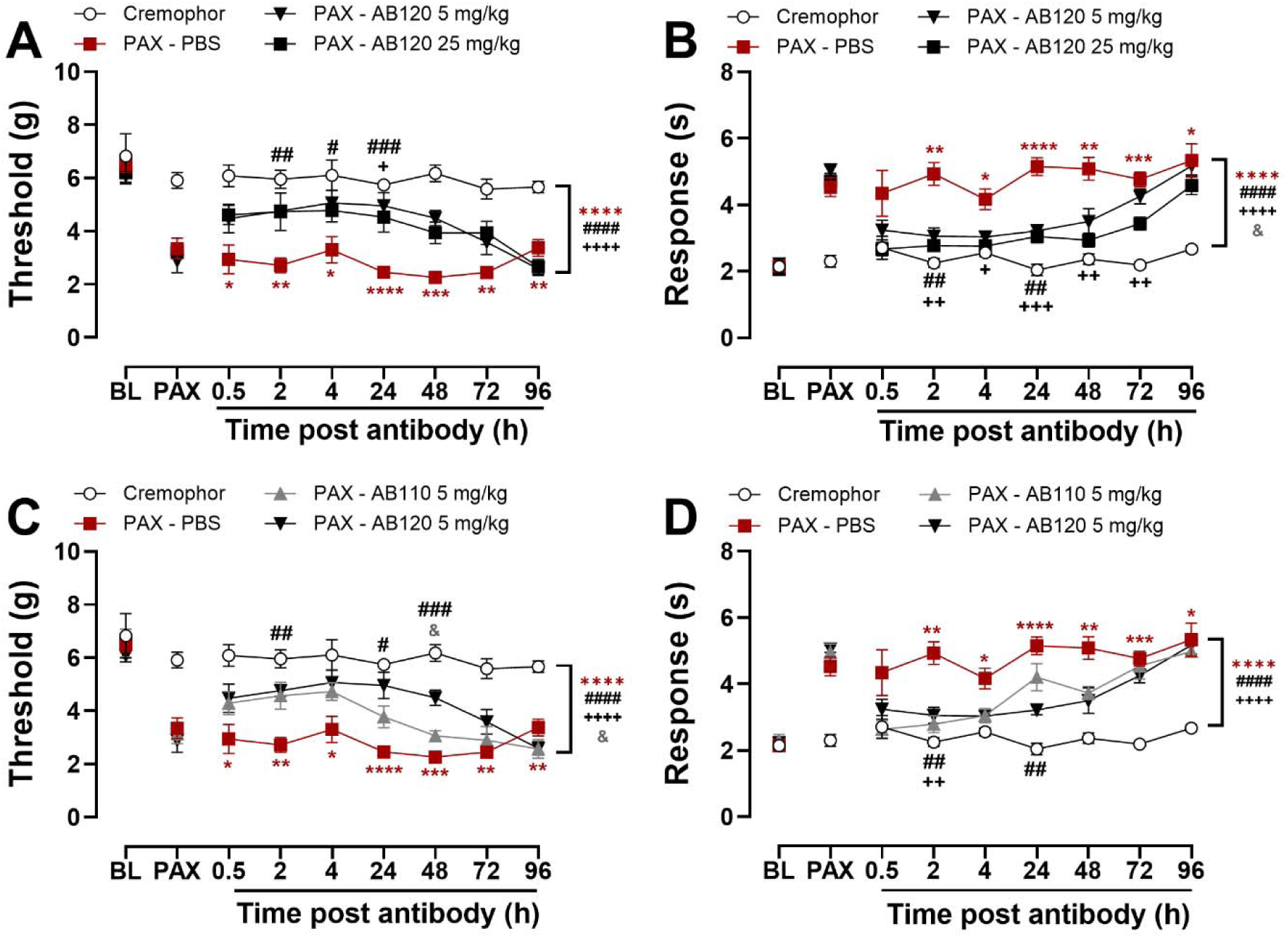
CB2-specific antibody agonist AB120 attenuates paclitaxel-induced mechanical and cold hypersensitivity in female mice. AB120 or AB110 was administered acutely at 14 days post initial paclitaxel dosing, when CIPN was established. AB120 (5 and 25 mg/kg, i.p.) attenuated paclitaxel-induced (**A**) mechanical and (**B**) cold hypersensitivity in mice. AB120 (5 mg/kg, i.p.) attenuated (**C**) mechanical hypersensitivity relative to AB110 (5 mg/kg, i.p.) or PBS treatment whereas both antibodies attenuated (**D**) cold hypersensitivity with comparable efficacy. Data show mean (± SEM) (n=4-6). Two-way ANOVA followed by Bonferroni’s *post hoc* test. Interaction between time points and treatment effect (A-B): *\*P<0.05 **P<0.01, ***P<0.001, ****P<0.0001 Cremophor vs. PAX-PBS; ^#^P <0.05, ^##^P <0.01, ^###^P <0.001 PAX-PBS vs. PAX-AB120 5 mg/kg; ^++^P <0.01, ^+++^P<0.001 PAX-PBS vs. AB120 25 mg/kg.* Interaction between time and treatment (C-D): *\*P<0.05, **P <0.01, ***P<0.001, ****P<0.0001 Cremophor vs. PAX-PBS; ^#^P<0.05, ^##^P<0.01, ^###^P <0.001 PAX-PBS vs. PAX-AB120 5 mg/kg; ^++^P<0.01 PAX-PBS vs. AB110 5 mg/kg. ^&^P<0.05, PAX-AB110 5 mg/kg vs. PAX-AB120 5 mg/kg. Brackets (main treatment effect): ****P<0.0001 Cremophor vs. PAX-treated groups; Panels A-B: ^####^P<0.0001 PAX-PBS vs. PAX-AB120 5 mg/kg; ^++++^P<0.0001 PAX-PBS vs. PAX-AB120 25 mg/kg; ^&^P<0.05 PAX-AB120 5 mg/kg vs. PAX-AB120 25 mg/kg; Panels C-D: ^####^P<0.0001 PAX-PBS vs. PAX-AB120 5 mg/kg; ^++++^P<0.0001 PAX-PBS vs. PAX-AB110 5 mg/kg; ^&^P<0.05 PAX-AB110 vs. PAX-AB120*.

AB120 (5 and 25 mg/kg, i.p.) attenuated paclitaxel-induced cold hypersensitivity in a treatment and time-dependent manner and cold responsiveness also changed across time (**Figure 5B**: Treatment: F_3,18_=34.62, p<0.0001; Time: F_3.495,62.92_=10.87, p<0.0001; Interaction: F_10.49,62.92_=2.359, p=0.0179). Overall, AB120 attenuated paclitaxel-induced mechanical hypersensitivity at both tested doses (5 mg/kg and 25 mg/kg: p<0.0001), and the high dose produced a greater suppression of mechanical hypersensitivity compared to the low dose (p=0.0172). Paclitaxel-treated mice developed marked cold hypersensitivity, as indicated by increased hind paw responses to acetone compared to PBS-treated controls at multiple time points: (2-96 h: p≤0.0159). AB120 (5 mg/kg, i.p.) reduced paclitaxel-induced cold hypersensitivity at 2 (p=0.0014) and 24 hours (p=0.0499) post-injection. AB120 (25 mg/kg, i.p.) attenuated paclitaxel-induced cold hypersensitivity from 2-72 h (p≤0.0435) post-injection.

Comparative analysis between AB120 and AB110 revealed differences in efficacy overall and across time (**Figure 5C**: Treatment: F_3,18_=45.06, p<0.0001; Time: F_3.343,60.17_=5.961, p=0.0008; Interaction: F_10.03,60.17_=2.291, p=0.0236). Both AB120 and AB110 reduced paclitaxel-induced mechanical hypersensitivity (p<0.0001 vs. Cremophor-vehicle) compared to the PBS-treated group (AB120: p<0.0001; AB110: p<0.0001), with AB120 producing a greater elevation of mechanical hypersensitivity compared to AB110 (p=0.0276) overall.

Paclitaxel reduced mechanical paw withdrawal thresholds compared to the PBS-treated group at all time points assessed (0.5-96 h: p≤0.0499). AB120 (5 mg/kg) attenuated paclitaxel-induced mechanical hypersensitivity at 2 (p=0.0013), 24 (p=0.0143), and 48 (p=0.0010) hours post-injection, whereas AB110 did not produce a reliable anti-allodynic effect in this experimental cohort. AB120 produced a greater elevation of paw withdrawal thresholds compared to AB110 at the 48-hour time point (p=0.0179).

Both AB110 and AB120 attenuated paclitaxel-induced cold hypersensitivity overall and in a time-dependent manner (**Figure 5D**: Treatment: F_3,18_=24.23, p<0.0001; Time: F_3.875,69.74_=12.27, p<0.0001; Interaction: F_11.62,69.74_=3.293, p=0.0009). Across the study duration, paclitaxel-treated mice consistently exhibited greater responses to acetone than Cremophor vehicle-treated controls (p<0.0001). Both AB120 and AB110 reduced paclitaxel-induced cold hypersensitivity across the observation period compared to PBS-treated mice (p<0.0001). Cold hypersensitivity was consistently observed in paclitaxel-treated mice, as evidenced by significantly increased responses to acetone compared to the Cremophor vehicle-treated control group from 2-96 hours post-treatment (p≤0.0153 for each timepoint). AB120 (5 mg/kg) reduced cold hypersensitivity at 2 (p=0.0091) and 24 hours (p=0.0017) post-injection. AB110 (5 mg/kg) also attenuated paclitaxel-induced cold hypersensitivity, but only at the 2-hour time point (p=0.0039).

### 3.6 CB2-specific antibody agonists AB110 and AB120 attenuate paclitaxel-induced allodynia without development of analgesic tolerance

In paclitaxel-treated mice, chronic administration of AB120 increased paw withdrawal thresholds overall but the interaction between treatment and time was not significant (**Figure 6A**: Treatment: F_3,18_=112.2, p<0.0001; Time: F_2.272,40.89_=0.2305, p=0.8218; Interaction: F_6.815,40.89_=0.2505, p=0.9671). Throughout the study, paclitaxel-treated mice displayed lower mechanical paw withdrawal thresholds compared to Cremophor vehicle-treated controls (p<0.0001). Chronic dosing with AB120 at both 5 and 25 mg/kg attenuated paclitaxel-induced mechanical hypersensitivity across the observation interval (5 mg/kg: p<0.0001; 25 mg/kg: p<0.0001), with no difference in efficacy observed between the two doses. This suggests that repeated administration of AB120 does not lead to analgesic tolerance under these experimental conditions.

**Figure 6.**
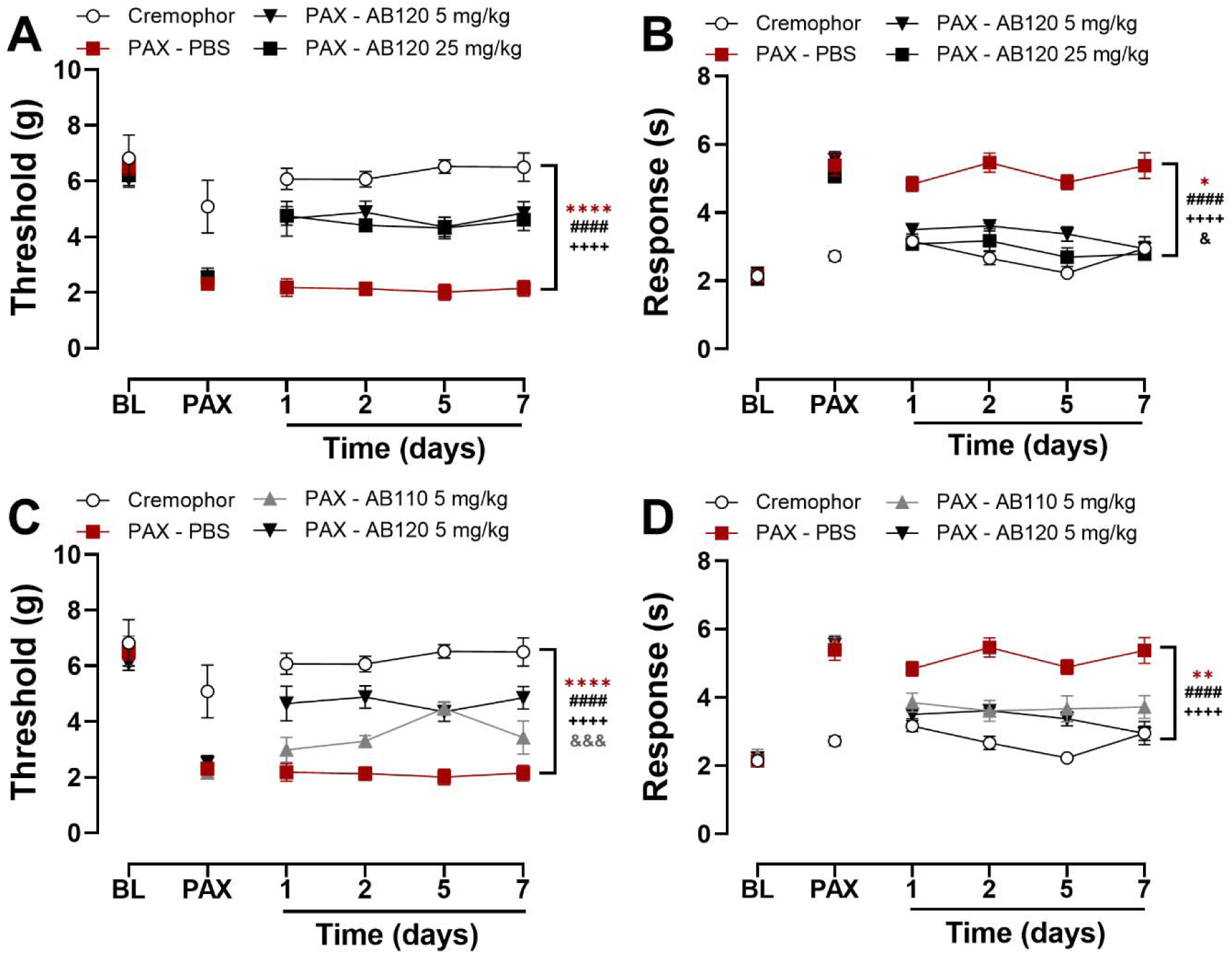
Tolerance did not develop to repeated daily systemic dosing with CB2-specific antibody agonists AB110 and AB120 in paclitaxel-treated female mice. AB120 (5 and 25 mg/kg, i.p.) produced sustained attenuation of paclitaxel-induced (**A**) mechanical and (**B**) cold hypersensitivity with repeated administration. AB120 5 mg/kg produced sustained attenuation of (**C**) mechanical hypersensitivity with superior efficacy compared to AB110 (5 mg/kg, i.p.). Both antibody agonists produced comparable suppression of paclitaxel-induced (**D**) cold hypersensitivity following repeated administration. Data show mean (± SEM) (n=4-6). Two-way ANOVA followed by Bonferroni’s *post hoc* test. *Brackets (main treatment effect): *P<0.05,**P<0.01,****P<0.0001 Cremophor vs. PAX-treated groups; Panels A-B: ^####^P<0.0001 PAX-PBS vs. PAX-AB120 5 mg/kg; ^++++^P<0.0001 PAX-PBS vs. PAX-AB120 25 mg/kg; ^&^P<0.05 PAX-AB120 5 mg/kg vs. PAX-AB120 25 mg/kg. Panel B: Cremophor vehicle did not differ from PAX-AB120 25 mg/kg; Panels C-D: ^####^P<0.0001 PAX-PBS vs. PAX-AB120 5 mg/kg; ^++++^P<0.0001 PAX-PBS vs. PAX-AB110 5 mg/kg; ^&&&^ P <0.001 PAX-AB110 vs. PAX-AB120*.

Repeated dosing with AB120 attenuated responses to hind paw acetone application overall in paclitaxel-treated mice and the interaction between time and treatment was not significant (**Figure 6B**: Treatment: F_3,18_=90.50, p<0.0001; Time: F_2.354,42.37_=2.639, p=0.0747; Interaction: F_7.062,42.37_=1.601, p=0.1609). Paclitaxel induced sustained cold hypersensitivity across the study duration, as evidenced by increased acetone responses compared to the Cremophor vehicle-treated group (p<0.0001). Both 5 and 25 mg/kg doses of AB120 attenuated paclitaxel-induced cold hypersensitivity following repeated administration, relative to PBS-treated mice (p<0.0001). The high dose of AB120 produced a greater suppression of paclitaxel-induced cold hypersensitivity compared to the low dose (p<0.0001).

Repeated daily dosing with AB120 (5 mg/kg, i.p.) and AB110 (5 mg/kg, i.p.) increased paw withdrawal thresholds in paclitaxel-treated mice across the observation interval (**Figure 6C**: Treatment: F_3,18_=71.05, p<0.0001; Time: F_2.613,47.03_=0.6635, p=0.5586; Interaction: F_7.838,47.03_=1.050, p=0.4129). As expected, paclitaxel-treated mice exhibited lower paw withdrawal thresholds compared to Cremophor vehicle-treated controls (p<0.0001) throughout the study. In paclitaxel-treated mice, both AB120 (5 mg/kg, i.p.) and AB110 (5 mg/kg, i.p.) attenuated mechanical hypersensitivity compared to PBS-treated controls (*p*<0.0001); however, the anti-allodynic efficacy of AB120 was greater than AB110 (*p*=0.0003), suggesting distinct therapeutic profiles.

Repeated dosing with AB120 (5 mg/kg, i.p.) and AB110 (5 mg/kg, i.p.) reduced time spent attending to acetone across the observation interval (**Figure 6D**: Treatment: F_3,18_=37.15, p<0.0001; Time: F_2.587,46.74_=1.308, p=0.2827; Interaction: F_7.790,46.74_=1.583, p=0.1574). As expected, paclitaxel-treated mice exhibited consistently elevated responses to acetone throughout the study compared to Cremophor vehicle-treated controls (p<0.0001). CB2-specific antibodies AB120 (5 mg/kg, i.p.) and AB110 (5 mg/kg, i.p.) attenuated paclitaxel-induced hypersensitivity to cold relative to PBS-treated mice (p<0.0001), with equivalent efficacy (p=0.3468).

### 3.7 Prophylactic dosing with AB120 attenuated, but did not prevent, paclitaxel-induced allodynia in mice

Prophylactic administration of AB120 (5 mg/kg, i.p.) increased paw withdrawal thresholds throughout the dosing interval overall relative to baseline (**Figure 7A**; baseline (BL) – day 8: Treatment: F_3,22_=13.82, p<0.0001; Time: F_3,66_=3.881, p=0.0128; Interaction: F_7.790,46.74_=1.583, p=0.1574). Paclitaxel treatment reduced paw withdrawal thresholds compared to Cremophor vehicle-treated mice (BL–day 8: p<0.0001). Prophylactic AB120 administration attenuated the development of paclitaxel-induced mechanical hypersensitivity compared to PBS-treated controls across the observation interval (p<0.0102). Prophylactic administration of AB120 (5 mg/kg, i.p.) decreased paclitaxel-induced cold hypersensitivity overall and in a time-dependent manner (**Figure 7B**; BL – day 8: Treatment: F_3,22_=47.71, p<0.0001; Time: F_3,66_=2.279, p=0.0876; Interaction: F_9,66_=2.055, p=0.0466). Paclitaxel produced persistent cold hypersensitivity, as indicated by elevated responses to acetone compared to Cremophor-treated mice (Day 2-8: p≤0.0006). Prophylactic AB120 (5 or 25 mg/kg, i.p.) attenuated the development of paclitaxel-induced cold hypersensitivity at early time points (Day 2-4: p≤0.0467 for each dose). Across the full prophylactic phase (BL–Day 8), both doses of AB120 reduced paclitaxel-induced cold hypersensitivity relative to PBS treatment (5 mg/kg: p=0.0002; 25 mg/kg: p<0.0001). However, this therapeutic effect was lost following treatment cessation (Days 11–15: p>0.99).

**Figure 7.**
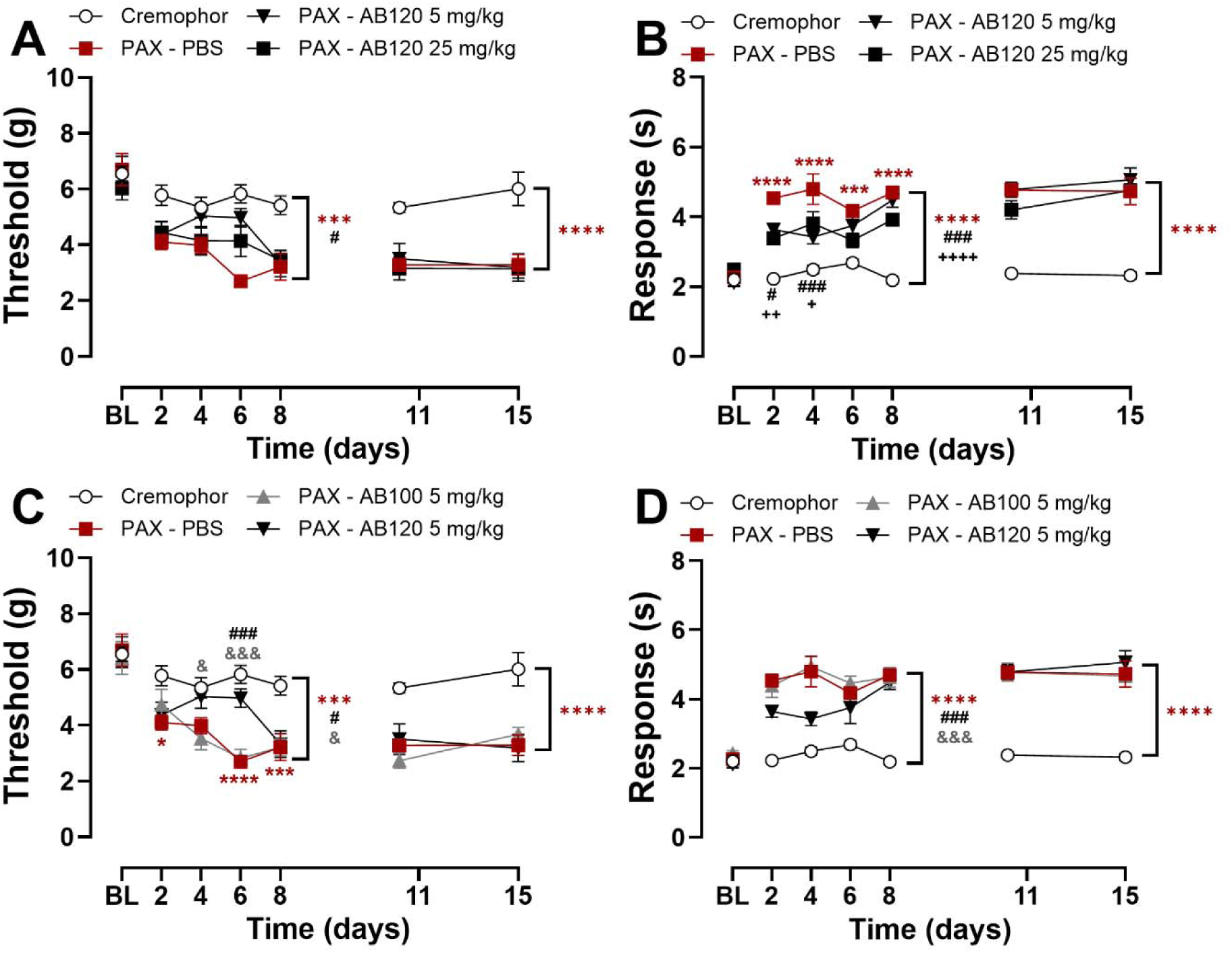
Prophylactic treatment with CB2-specific antibody agonist AB120 delays but does not prevent development of paclitaxel-induced mechanical and cold hypersensitivity in female mice. Repeated prophylactic dosing with AB120 (5 or 25 mg/kg, i.p.) suppressed the development of paclitaxel-induced (**A**) mechanical and (**B**) cold hypersensitivity. Inactive isotype control AB100 did not alter the development of paclitaxel-induced (**C**) mechanical and (**D**) cold hypersensitivity. Data show mean (± SEM) (n=6). Two-way ANOVA followed by Bonferroni’s *post hoc* test. Interaction between time points and treatment effect (A-B): *\*\*\*P<0.001, ****P<0.0001 Cremophor vs. PAX-PBS; ^#^P<0.05, ^###^P<0.001 PAX-PBS vs. PAX-AB120 5 mg/kg; ^++^P<0.05, ^++^P<0.01 PAX-PBS vs. AB120 25 mg/kg.* Interaction between time and treatment (C-D): *\*P<0.05, ***P<0.001, ****P<0.0001 Cremophor vs. PAX-PBS; ^###^P<0.001 PAX-PBS vs. PAX-AB120 5 mg/kg; ^&^P<0.05, ^&&&^P<0.001 PAX-AB100 5 mg/kg vs. PAX-AB120 5 mg/kg. Brackets (main treatment effect): **P<0.01,***P<0.001, ****P<0.0001 Cremophor vs. PAX-treated groups; Panels A-B: ^#^P<0.05, ^####^P<0.0001 PAX-PBS vs. PAX-AB120 5 mg/kg; ^++++^P<0.0001 PAX-PBS vs. PAX-AB120 25 mg/kg; Panels C-D: ^#^P<0.05, ^###^P<0.001 PAX-PBS vs. PAX-AB120; ^&^P<0.05, ^&&&^P <0.001 PAX-AB100 vs. PAX-AB120*.

Prophylactic AB120 (5 mg/kg, i.p.) treatment attenuated development of paclitaxel-induced mechanical hypersensitivity compared to treatment with PBS or the isotype control AB100 (5 mg/kg, i.p.) during the dosing regimen (**Figure 7C**; BL – day 8: Treatment: F_3,21_=18.18, p<0.0001; Time: F_3,63_=5.432, p=0.0022; Interaction: F_9,63_=2.867, p=0.0067). Paclitaxel reduced mechanical thresholds on days 2, 6, and 8 of prophylactic treatment compared to Cremophor vehicle-treated controls (p≤0.0208 for each timepoint). AB120 (5 mg/kg, i.p.) attenuated mechanical hypersensitivity by day 6 compared to PBS-treated mice (p=0.0003) and showed greater efficacy than the isotype control AB100 on days 4 (p=0.0380) and 6 (p=0.0008) post-treatment. Throughout the observation period, paclitaxel-treated mice receiving PBS prophylactically exhibited robust mechanical hypersensitivity relative to Cremophor vehicle-treated controls (BL – day 8: p<0.0001). AB120 delayed the onset of paclitaxel-induced mechanical hypersensitivity compared to PBS (p=0.0051) or the isotype control AB100 (p=0.0124).

Prophylactic AB120 (5 mg/kg, i.p.) attenuated development of paclitaxel-induced cold hypersensitivity compared to treatment with PBS or the isotype control AB100 (5 mg/kg, i.p.) across the treatment regimen (**Figure 7D**; BL–day 8: Treatment: F_3,21_=71.44, p<0.0001; Time: F_3,63_=1.194, p=0.3194; Interaction: F_9,63_=1.789, p=0.0880). Overall, paclitaxel-treated mice receiving PBS exhibited elevated acetone responses compared to Cremophor vehicle-treated controls throughout the study (p<0.0001). Prophylactic treatment with AB120 (5 mg/kg, i.p.) attenuated cold hypersensitivity overall relative to PBS-treated mice (p=0.0004) or the inactive isotype control AB100 (p=0.0001). However, the therapeutic benefit of prophylactic AB120 was lost following treatment cessation during the washout period (Day 11 to Day 15: p>0.99).

Paclitaxel treatment reliably reduced paw withdrawal thresholds and increased response to acetone compared to Cremophor-vehicle-treated mice after treatment with AB120 was discontinued in all treatment regimen comparisons (**Figure 7A**; day 11 – 15: Treatment: F_3,22_=16.96, p<0.0001; Time: F_1,22_=0.0822, p=0.7770; Interaction: F_3,22_=0.4208, p=0.7399; **Figure 7B**; day 11 – 15: Treatment: F_3,22_=45.67, p<0.0001; Time: F_1,22_=1.089, p=0.3081; Interaction: F_3,22_=0.7338, p=0.5430; **Figure 7C**; day 11 – 15: Treatment: F_3,21_=18.51, p<0.0001; Time: F_1,21_=1.161, p=0.2934; Interaction: F_3,21_=0.9332, p=0.4422; **Figure 7D**; day 11 – day 15: Treatment: F_3,21_=48.94, p<0.0001; Time: F_1,21_=0.0056, p=0.9407; Interaction: F_3,21_=0.3048, p=0.8216).

### 3.8 Anti-allodynic efficacy of AB120 is absent in global CB2 knock out mice

We investigated whether ability of AB120 to suppress the maintenance of paclitaxel-induced mechanical hypersensitivity depends on CB2 receptors by using CB2KO mice. ABT120 (5 mg/kg, i.p.) suppressed paclitaxel-induced mechanical hypersensitivity in WT but not CB2 KO mice (**Figure 8A**: Treatment: F_2,12_=10.70, p=0.0022; Time: F_4.082,48.98_=5.588, p=0.0008; Interaction: F_8.163,48.98_=4.909, p=0.0002); this suppressive effect was observed across the observation interval (*p*<0.0001) and in a time-dependent manner. In WT mice, AB120 (5 mg/kg) reduced paclitaxel-induced mechanical hypersensitivity for at least 24 hours post-treatment (2-24 h: *p*=≤0.0443) compared to CB2KO mice receiving the same treatment. In CB2KO mice, AB120 showed no significant difference from PBS at any time point (*p*>0.99).

**Figure 8.**
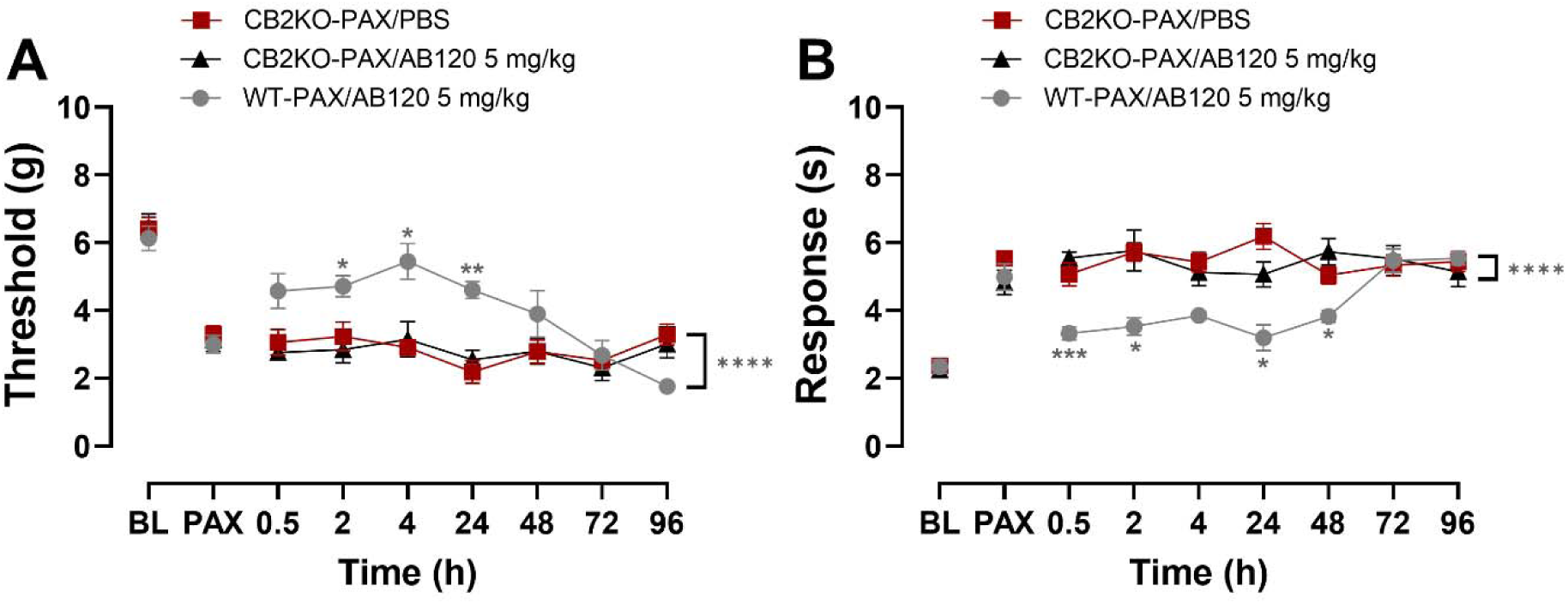
Analgesic efficacy of CB2-specific antibody AB120 is absent in global CB2 knock out mice (CB2KO). AB120 (5 mg/kg, i.p.) attenuated (**A**) mechanical and (**B**) cold hypersensitivity in female wild type (WT), but not CB2KO mice. Data show mean (± SEM) (n=5). Two-way ANOVA followed by Bonferroni’s *post hoc* test. Interaction between time points and treatment effect: *\*P<0.05, **P<0.01, ***P<0.001 WT-PAX/AB120 vs. CB2KO-PAX/AB120. Brackets (main treatment effect): ****P<0.0001 WT-PAX/AB120 vs. CB2KO-PAX/AB120*.

ABT120 (5 mg/kg, i.p.) similarly suppressed paclitaxel-induced cold hypersensitivity in WT but not CB2 KO mice (**Figure 8B**: Treatment: F_2,12_=21.07, p=0.0001; Time: F_3.520,42.24_=2.907, p=0.0382; Interaction: F_7.041,42.24_=4.942, p=0.0004). Overall, AB120 (5 mg/kg, i.p.) effectively reduced cold hypersensitivity in wild type but not in CB2 KO mice across the observation interval (*p* < 0.0001). In wild-type mice, AB120 (5 mg/kg) attenuated paclitaxel-induced cold hypersensitivity over 48-hours post-treatment (0.5 h: *p*=0.0001; 2 h: *p*=0.0494; 24 h: *p*=0.0247; 48 h: *p*=0.0147). In contrast, acetone responses did not differ between PBS- and AB120-treated CB2KO mice at any time point.

### 3.9 AB120, a CB2-specific antibody agonist, reduces paclitaxel-induced hypersensitivity in mammary tumor-bearing mice

BALB/c female mice were orthotopically injected with 4T1 mammary carcinoma cells into the second mammary fat pad to ascertain whether the anti-allodynic effects of AB120 were preserved in tumor-bearing mice. Once tumors became palpable, tumor growth was monitored daily using calipers, and the paclitaxel dosing regimen (10 mg/kg, i.p. on 3 alternate days) was initiated (**Figure 9A**). Tumor volume increased markedly over a 7-day monitoring period, confirming successful establishment of the tumor-bearing mouse model (**Figure 9B)**.

**Figure 9.**
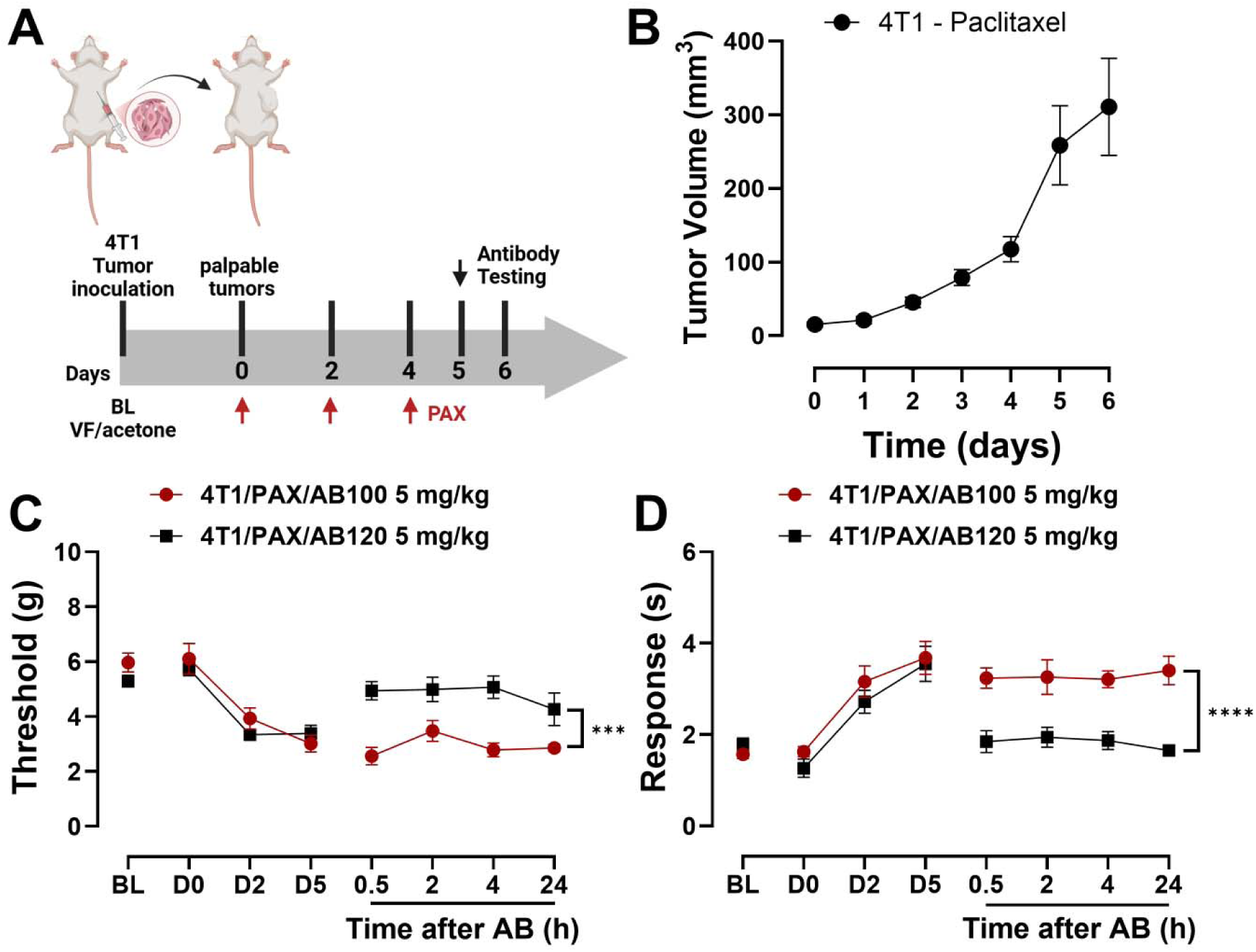
CB2-specific antibody agonist AB120 attenuates paclitaxel-induced mechanical and cold hypersensitivity in mammary tumor bearing mice. (**A**) Timeline of tumor inoculation and antibody efficacy study. Injection of 4T1 cells into the mammary fat pad produced exponential growth evidenced by (**B**) tumor volume (mm^3^). Treatment with paclitaxel (10 mg/kg i.p. on 3 alternate days) produced robust (**C**) mechanical and (**D**) cold hypersensitivity in tumor-bearing mice. AB120 (5 mg/kg, i.p.) attenuated paclitaxel-induced mechanical and cold hypersensitivity in tumor-bearing mice, whereas inactive isotype control AB100 (5 mg/kg. i.p.) did not alter responses to cutaneous stimulation. Data show mean (± SEM) (n=6). Two-way ANOVA followed by Bonferroni’s *post hoc* test. Interaction between time points and treatment effect: *Brackets (main treatment effect): ***P<0.001, ****P<0.0001 4T1/PAX/AB120 5 mg/kg vs. 4T1/PAX/AB100 5 mg/kg*.

Mechanical thresholds and acetone responses did not change in experimental groups between baseline (BL) and time when mammary tumors became palpable and paclitaxel regimen was initiated (day 0, D0) (BL vs. D0: **Figure 9C**: AB100: t=0.2547, df=5, p=0.8091; AB120: t=2.000, df=5, p=0.1020; **Figure 9D**: AB100: t=0.4053, df=5, p=0.6999; AB120: t=2.520, df=5, p=0.0532). Paclitaxel decreased mechanical paw withdrawal thresholds in tumor-bearing mice. Acute treatment with AB120 (5 mg/kg, i.p. on day 5 (D5)) increased mechanical paw withdrawal thresholds in paclitaxel-treated tumor bearing mice overall relative to isotype control AB100 (5 mg/kg, i.p.) but the interaction between treatment and time was not significant (**Figure 9C**: Treatment: F_1,10_=25.95, p=0.0005; Time: F_2.039,20.39_=1.594, p=0.2272; Interaction: F_2.039,20.39_=1.276, p=0.3012). Over the 24-hour observation period, AB120 increased paw withdrawal thresholds compared to AB100 (p=0.0005), demonstrating its anti-allodynic efficacy in a paclitaxel-induced neuropathic pain model in tumor-bearing mice.

Paclitaxel treatment increased responsiveness to acetone in tumor-bearing mice, consistent with development of cold allodynia. AB120 (5 mg/kg, i.p.) reduced acetone responses in paclitaxel-treated, tumor-bearing mice (**Figure 9D**: Treatment: F_1,10_=44.31, p<0.0001; Time: F_1.831,18.31_=0.0439, p=0.9468; Interaction: F_1.831,18.31_=0.4533, p=0.6256) overall compared to AB100 (p<0.0001). Thus, AB120 attenuated paclitaxel-induced mechanical and cold hypersensitivity in tumor-bearing mice.

### 3.10 CB2-specific antibody agonists do not produce motor ataxia or locomotor deficit in mice

No reliable differences were observed between treatment groups in the rota-rod test on the test day or relative to their respective baseline performance (**Figure 10A**: Treatment: F_5,27_=1.012, p=0.4297; Time: F_1,27_=1.460, p=0.2373; Interaction: F_5,27_=1.837, p=0.1392), indicating that none of the treatments impaired motor coordination at this time-point. CP55,940 decreased descent latency relative to PBS treatment (t=2.303 df=8, p=0.051, one-tailed unpaired t-test) and relative to pre-injection baseline performance (t=2.44 df=4, p=0.0044), paired t-test).

**Figure 10.**
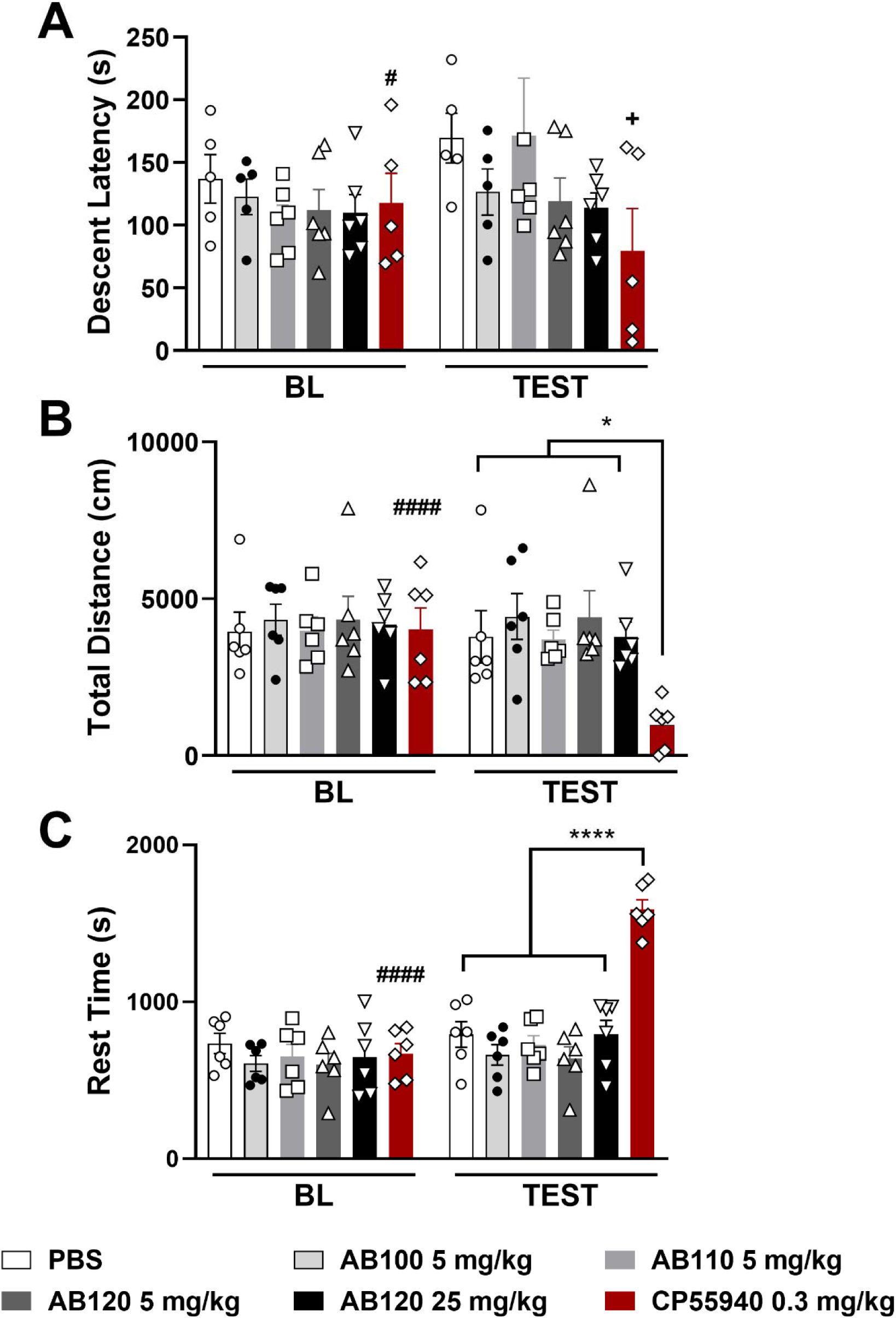
Systemic treatment with CB2-specific antibody agonists does not produce motor ataxia or locomotor deficits in naïve mice. AB100 (5 mg/kg), AB110 (5 mg/kg), AB120 (5 or 25 mg/kg) i.p. did not reduce (**A**) descent latency in the rota-rod test. CP55,940 (0.3 mg/kg, i.p.) reduced descent latency relative to either vehicle injection (p=0.0251, unpaired t-test, one-tailed) or baseline pre-injection performance (p=0.0441, paired t-test, one-tailed). CP55-940 (0.3 mg/kg, i.p.), but not CB2 antibody agonists (**B**) decreased total distance traveled and (**C**) increased rest time performed in the activity meter test. Locomotor behaviors were assessed 2 hours following all treatments over a 30-min observation interval. Data show mean (± SEM) (n=5-6). Two-way ANOVA followed by Bonferroni’s *post hoc* test. *Brackets: *P <0.05, ****P <0.0001 CP55-940 0.3 mg/kg vs all other groups. Symbols on top of bars: ^#^P<0.05, ^####^P<0.0001 BL vs. test; ^+^P<0.05 vs. control PBS*.

Small molecule cannabinoid agonist CP55,940, but not antibody agonists, reduced total distanced traveled in the activity meter post-injection, as revealed by a significant interaction between time and treatment (**Figure 10B**: Treatment: F_5,30_=1.520, p=0.2134; Time: F_1,30_=10.99, p=0.0024; Interaction: F_5,30_=7.015, p=0.0002). CP55,940 (0.3 mg/kg, i.p.) reduced the total distance traveled compared to all other treatment groups on test day (p≤0.0338 for each comparison). CP55,940 also reduced total distance traveled compared to its baseline (pre-injection) performance (p<0.0001). Thus, CB2-targeting antibodies do not impair locomotor function, unlike the non-selective cannabinoid agonist CP55,940.

CP55,940, but not antibody agonists, also increased rest time in the activity meter overall and the interaction between time and treatment was significant (**Figure 10C**: Treatment: F_5,30_=8.071, p<0.0001; Time: F_1,30_=83.30, p<0.0001; Interaction: F_5,30_=36.35, p<0.0001). Specifically, CP55,940 increased rest time compared to all other treatment groups (p<0.0001 for each comparison). Moreover, CP55,940 reduced locomotor activity relative to baseline (p<0.0001), highlighting that only the synthetic agonist, and not the therapeutic antibodies, impaired movement in this assay.

### 3.11 Paclitaxel-induced cytotoxicity in 4T1 cells is preserved in the presence of CB2-specific antibody agonists

We assessed the impact of CB2-specific antibody agonists and paclitaxel over a wide range of molar concentrations on 4T1 and HEK293 cell viability (**Supplemental Table 3**). As expected, paclitaxel (EC50=40 nM; **Figure 11A**) reduced 4T1 tumor cell viability (%Change), whereas AB120 did not reduce 4T1 cell viability (EC95=49.359 µM, **Figure 11B**). The impact of various concentrations of AB120 on the dose response curve of paclitaxel in suppression of 4T1 tumor cell viability is shown in **Figure 11C**. Similarly, **Figure 11D** shows the effect of various concentrations of paclitaxel on the dose response curve of AB120 on 4T1 tumor cell viability. Combination of paclitaxel and AB120 produced a likely additive effect in suppression of 4T1 tumor cell viability, as observed in synergy scores (**Supplemental Table 3**). Computational quantification of the drug combination response, plotted as a three-dimensional synergy map over the dose matrix, indicated that the combination of AB120 and paclitaxel on 4T1 cells was additive using the Bliss (**Figure 11E**) and HSA models (**Figure 11F**), whereas the LOEWE model could not calculate a synergy score for that specific data set (**Figure 11G**).

**Figure 11.**
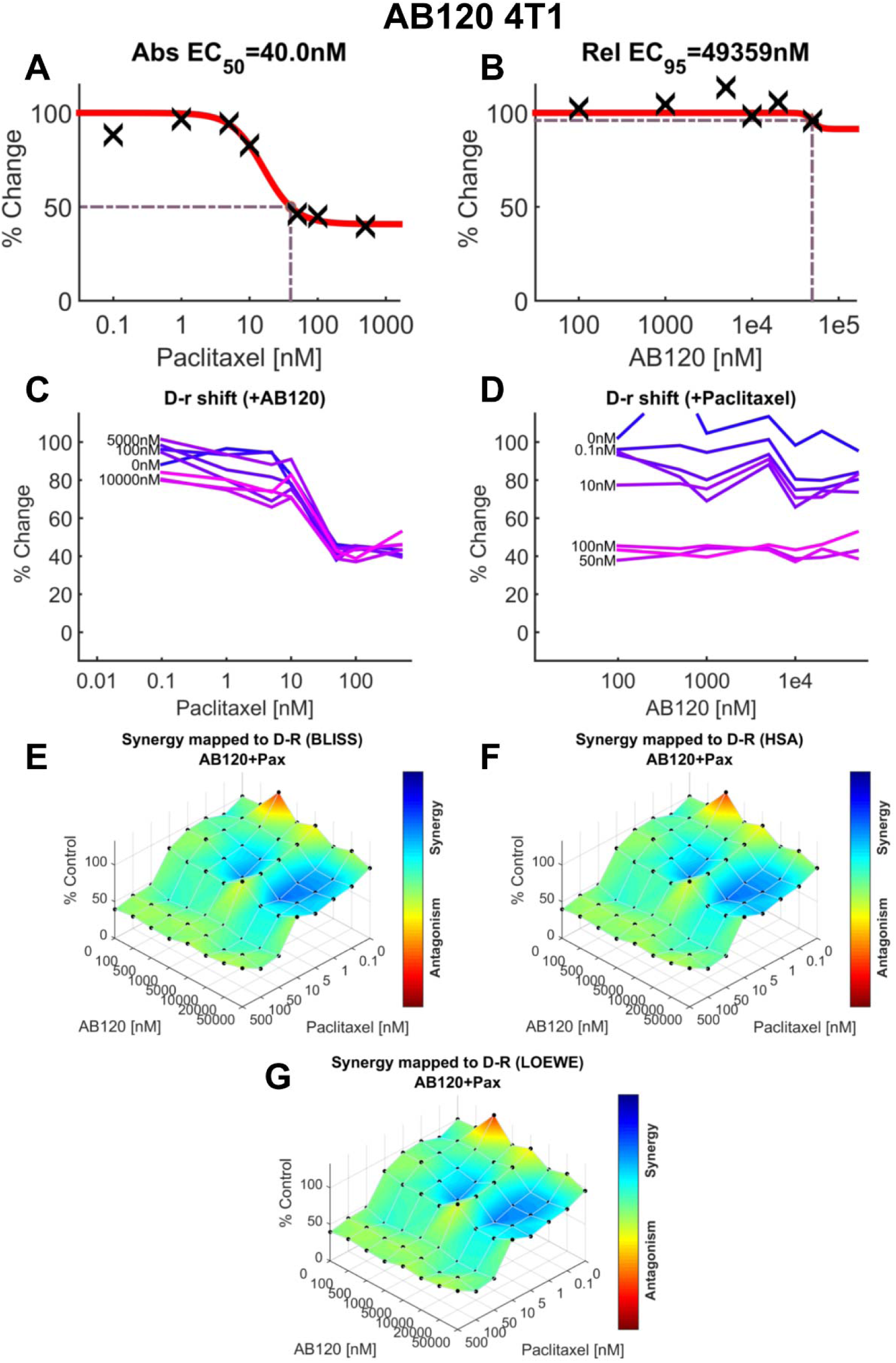
AB120 produces additive effects with paclitaxel in 4T1 cells to reduce murine mammary carcinoma (4T1) cell line viability. (**A**) Dose-response for the effect of paclitaxel in 4T1 cell viability (EC50=40 nM). (**B**) Dose-response for the effect of AB120 in 4T1 cells (EC95=49.359 µM). (**C**) Paclitaxel dose response shift observed in the presence of increasing concentrations of AB120. (**D**) AB120 dose response shift observed in the presence of increasing concentrations of paclitaxel. (**E–G**) The 3-dimensional landscape of the dose matrix is represented on a color scale, where blue reflects evidence of synergy and red reflects evidence of antagonism. The landscape of the dose matrix of combination responses for AB120 and paclitaxel based on the (**E**) Bliss model, (**F**) Highest Single Agent (HSA) model and (**G**) Loewe model. Each model supports dose dependent additive effects of the combination in reducing 4T1 tumor cell line viability. Cell viability is plotted as % control. (n = 3 experiments).

In contrast, paclitaxel (EC95=13 nM, **Figure 12A**) showed limited efficacy on viability of non-tumor HEK293 cells. In addition, AB120 (EC95=233.362 µM) did not produce alterations in HEK293 cell viability (**Figure 12B**). The lack of effect of AB120 on various concentrations of paclitaxel is shown by the curve in **Figure 12C**. Similarly, the lack of effect of various concentrations of paclitaxel to the dose response of AB120 is shown in **Figure 12D**. Synergy scores demonstrate that combination of paclitaxel and AB120 produced only modest additive effects in HEK293 cell viability (**Supplemental Table 3**). The three-dimensional synergy maps plotted over a dose matrix revealed an additive effect of the combination of AB120 and paclitaxel according to the Bliss (**Figure 12E**), HSA (**Figure 12F**) and LOEWE models (**Figure 12G**).

**Figure 12.**
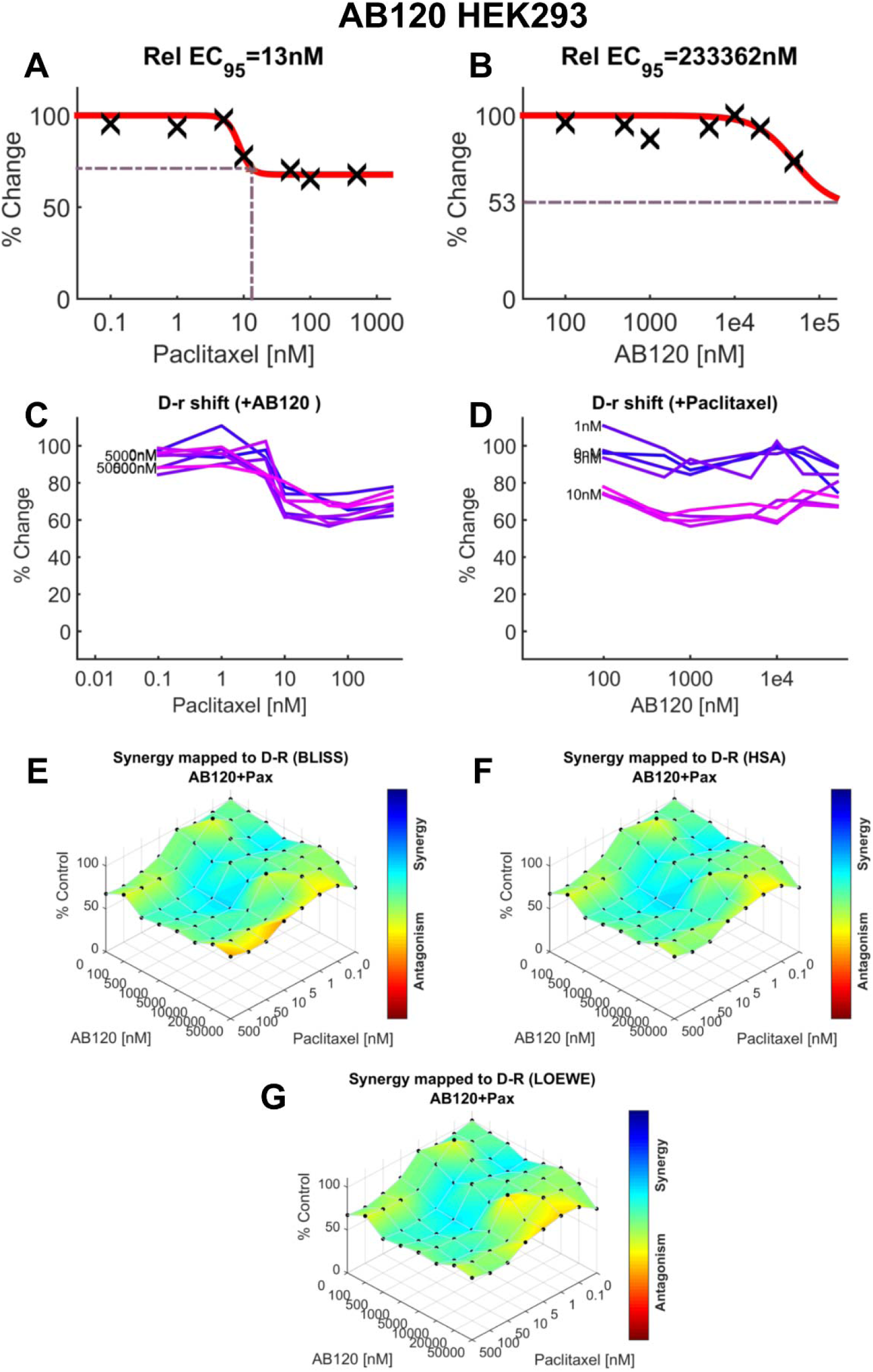
AB120 shows likely additive effect in the viability of non-tumor HEK293 cells in combination with paclitaxel. (**A**) Dose-response for the effect of paclitaxel on viability of HEK293 cells (EC95=13 nM). (**B**) Dose-response for the effect of AB120 on viability of HEK293 cells (EC95=233.362 µM). (**C**) Paclitaxel dose response shift observed in the presence of increasing concentrations of AB120 in HEK293 cell viability. (**D**) AB120 dose response shift observed in the presence of increasing concentrations of paclitaxel in HEK293 cell viability. (**E–G**) The 3-dimensional landscape of the dose matrix is represented on a color scale, where blue reflects evidence of synergy and red reflects evidence of antagonism. The landscape of the dose matrix combination responses for AB120 and paclitaxel based on the (**E**) Bliss model, (**F**) Highest Single Agent (HSA) model and (**G**) Loewe model. Cell viability is plotted as % control (n = 4 experiments).

As anticipated, paclitaxel reduced 4T1 tumor cell viability (EC50=21.5 nM; **Supplemental Figure 1A**), while AB110 showed minimal impact on viability (EC50 >50.000 µM; **Supplemental Figure 1B**). The influence of different concentrations of AB110 on paclitaxel’s dose-response curve for inhibition of 4T1 cell viability is presented in **Supplemental Figure 1C**. Likewise, **Supplemental Figure 1D** illustrates how varying doses of paclitaxel affect the dose-response curve of AB110. When combined, paclitaxel and AB110 produced an apparently additive reduction in 4T1 tumor cell viability, consistent with the synergy scores in **Supplemental Table 3**. Computational analysis of the combination response, shown as a three-dimensional synergy map across the dose matrix, demonstrated that the combination of paclitaxel and AB110 behaves additively according to the Bliss (**Supplemental Figure 1E**), HSA (**Supplemental Figure 1F**) and LOEWE models (**Supplemental Figure 1G**).

As expected, paclitaxel also decreased 4T1 tumor cell viability (EC50=31.9 nM, **Supplemental Figure 2A**) in the analysis with the isotype control AB100 (EC95=4.451 µM, **Supplemental Figure 2B**). **Supplemental Figure 2C** shows how varying concentrations of AB100 do not alter the paclitaxel dose-response curve in suppressing 4T1 cell viability. Similarly, **Supplemental Figure 2D** depicts the lack of effect of different paclitaxel doses on the AB100 dose-response curve. Combination of paclitaxel and AB100 produced what appears to be an additive reduction in 4T1 tumor cell viability, in agreement with the synergy scores provided in **Supplemental Table 3**. Three-dimensional synergy mapping across the dose matrix further confirmed additive interactions according to the Bliss (**Supplemental Figure 2E**) and HSA models (**Supplemental Figure 2F**), but a slight antagonism according to the LOEWE model (**Supplemental Figure 2G**).

Paclitaxel exhibited minimal impact on the viability of non-tumor HEK293 cells (EC50 = 123 nM; **Supplemental Figure 3A**). Likewise, AB110 (EC50> 50.000 µM) did not alter non-tumor cell viability (**Supplemental Figure 3B**). The absence of any effect from AB110 across different paclitaxel concentrations is illustrated in **Supplemental Figure 3C**, and **Supplemental Figure 3D** similarly shows that different paclitaxel doses did not alter AB110’s dose-response profile. Synergy scores indicate that the combination of paclitaxel and AB110 yielded additive effects on non-tumor cell viability (**Supplemental Table 3**). Consistently, three-dimensional synergy maps generated across the dose matrix showed additive interactions between AB110 and paclitaxel in HEK293 cells according to the Bliss (**Supplemental Figure 3E**), HSA (**Supplemental Figure 3F**), and LOEWE models (**Supplemental Figure 3G**).

Accordingly, paclitaxel (EC50>500 nM; **Supplemental Figure 4A**) and AB100 (EC95=17 nM; **Supplemental Figure 4B**) showed only limited effects on the viability of non-tumor HEK293 cells. **Supplemental Figure 4C** demonstrates that AB100 had no impact across varying paclitaxel concentrations, and **Supplemental Figure 4D** likewise shows that different doses of paclitaxel did not modify AB100’s dose–response behavior in non-tumor cells. Synergy scores indicate that the combination of paclitaxel and AB100 produced additive effects on HEK293 cell viability (**Supplemental Table 3**). In alignment with these findings, three-dimensional synergy maps across the dose matrix revealed additive interactions under the Bliss (**Supplemental Figure 4E**), HSA (**Supplemental Figure 4F**), and LOEWE models (**Supplemental Figure 4G**).

## 4 Discussion

No therapeutic antibody targeting GPCRs is clinically available to treat chronic neuropathic pain [33; 44; 62]. This is the first report of CB2 activation by a GPCR antibody agonist in a preclinical pain model and first demonstration of CB2-mediated analgesia in paclitaxel-treated tumor-bearing mice. Activation of CB2 receptors reduces pain-related behaviors in preclinical models without unwanted side effects [7; 15; 16; 20; 23; 37; 56-58]. However, clinical translation has been limited by rapid drug clearance, off-target effects, and inconsistent results in human trials [6; 9; 50; 66]. Therapeutic antibodies offer potential advantages to small molecules (e.g., limited CNS exposure, high target affinity and extended half-life) [8; 62]. Here, we show that CB2-specific therapeutic antibody agonists, AB110 and AB120, attenuate paclitaxel-induced mechanical and cold hypersensitivities in a mouse CIPN model. Notably, these antibodies did not produce tolerance or motor impairment. These observations align with recent studies from our group that reinforce a role for peripheral CB2 receptors in analgesic efficacy of CB2 agonists in pathological pain models [7; 22; 24; 38].

Therapeutic antibodies used in chronic conditions often have long durations of action due to their extended half-lives, with effects lasting weeks to months [61]. This allows for infrequent dosing schedules, typically ranging from biweekly to every few months, depending on the specific antibody and condition treated [63]. Despite significant efforts made towards developing monoclonal antibodies for chronic pain, only those targeting migraine have reached the market [49]. Here we demonstrate that CB2 receptor antibody agonists AB110 and AB120 present cAMP biased agonism in mouse macrophage cell lines. The decrease in cAMP production was blocked by a selective CB2 antagonist/inverse agonist SR144528, in accordance to our previous data in macrophage and human cell lines [21]. In addition, AB110 and AB120 recruit β-arrestin-2 with lower potency and efficacy, when compared to small molecule CB2 agonist HU-308. Considering receptor internalization and down regulation of CB2 receptors led by β-arrestin-2 [2; 10], the signaling bias of CB2 antibody agonists AB110 and AB120 could represent an advantage in cannabinoid therapeutics.

AB120 showed greater and longer-lasting effects than AB110, reducing pain behavior relative to controls (PBS, AB100) for up to 48 hours. Small molecule CB2 agonists (e.g. JWH133, AM1241) reduce pathological pain in animal models, but their effects are typically short-lived [16; 20; 37; 57; 58]. Efficacy of AM1710 largely diminishes within 24 hours in the absence of chronic treatment [15; 37]. Monoclonal antibodies exhibit peripherally restricted activity due to their large molecular size, which limits their ability to cross the blood-brain barrier and confines their effects to peripheral tissues [63]. Because CB2 activation in the periphery suppresses mRNA expression levels of pro-inflammatory cytokines [15; 24; 37], attenuates pain behavior [7; 22; 24; 37; 39], produces neuroprotection and circumvents CB1-associated psychotropic effects [15; 30], the target specificity and peripherally-restricted characteristic of antibodies targeting CB2 receptors may represent a safer option for pain modulation compared to conventional analgesic strategies. These CB2 activating antibodies also attenuate inflammatory markers associated with fibrosis, as well as inflammation and scarring in human liver tissue [21]. Accordingly, we demonstrate that AB110 and AB120 reduce the expression of LPS-induced pro-inflammatory cytokines and chemokines characteristic of M1 macrophage stage *in vitro*. However, the effect of these CB2 antibodies in IL-4 promoted anti-inflammatory markers (M2) was more varied with AB120 showing a broader effect for attenuation of all M2 macrophage markers, when compared to AB110. Thus, CB2-targeted antibodies attenuate inflammatory markers associated with neuropathic pain and may provide a longer-lasting alternative to small molecule CB2 agonists for managing painful symptoms in CIPN.

In our study, CB2-specific antibody candidates AB110 and AB120 did not produce analgesic tolerance in paclitaxel-treated mice under the present dosing conditions. Daily injections produced sustained suppression of paclitaxel-induced mechanical and cold hypersensitivity for up to seven consecutive days. AB120 showed superior efficacy in alleviating paclitaxel-induced mechanical hypersensitivity with repeated dosing compared to AB110. Using small molecule CB2 agonists, we observed sustained therapeutic efficacy that was absent in global KO [15] and cKO mice lacking CB2 in primary sensory neurons [7; 22; 24]. CB2 is also notably expressed on peripheral immune cells [3], and their activation may avoid psychoactive effects such as reward and tolerance. Unlike CB1 agonists and opioid analgesics, small-molecule CB2 agonists do not induce tolerance [15; 38]. In preclinical models, CB2 activation reduces neuropathic nociception, provides opioid-sparing effects, and decreases behaviors associated with opioid dependence, including conditioned place preference, self-administration, and withdrawal symptoms [7; 15; 22; 37; 38]. The absence of analgesic tolerance highlights the potential tolerability and safety of CB2-specific antibodies for treating chronic pain conditions such as CIPN. Longer-term studies are needed to assess potential risks associated with immunogenicity, particularly the development of anti-drug antibodies (ADAs). While the incidence of immunogenic responses has been relatively low in the treatment of painful conditions such as migraine [11], it remains a critical concern for biologic therapies, including monoclonal antibodies and fusion proteins, which can elicit immune reactions that may compromise their long-term efficacy or safety [73].

Prophylactic treatment with AB120, our most efficacious CB2-specific antibody agonist, delayed, but did not prevent, development of paclitaxel-induced mechanical and cold hypersensitivities; the protective effect diminished once treatment was discontinued. Chemotherapy agents can exert neurotoxic effects on both the peripheral and central nervous systems [73]. The condition is primarily sensory in nature, with symptoms including paresthesia, numbness, and pain, sometimes emerging after just a single dose of chemotherapy [29]. Even after treatment is discontinued, symptoms may persist for years or become permanent [18]. Treatment options for CIPN remain limited and most drugs used in clinical practice provide only symptomatic relief and do not prevent the onset of CIPN, largely due to the incomplete understanding of its underlying mechanisms [45]. Factors such as patient variability, challenges in predicting CIPN development, and inadequate assessment methods further hinder the effectiveness of prophylactic interventions [35]. Despite this shortcoming, small-molecule CB2 agonists can block or attenuate the development of paclitaxel-induced behavioral hypersensitivities in preclinical studies for weeks [22; 23; 54]. In line with our observations using larger antibody-based agonists, anti-allodynic effects of small-molecule CB2 agonists dissipate after treatment cessation, leading to the reemergence of CIPN-related sensory disturbances [22; 23]. Nonetheless, CB2-specfic antibody agonists may provide an alternative approach to achieve long lasting amelioration of CIPN symptom development with convenient dosing schedules. Further studies with longer treatment regimens are necessary to address current limitations.

While most studies assess CIPN in tumor-free mice, we additionally evaluated the ability of CB2-specific antibody agonists to suppress paclitaxel-induced behavioral hypersensitivities in mice bearing mammary tumors. This approach provides a more clinically relevant model that better reflects the human experience. Our data demonstrate that orthotopic inoculation of 4T1 cells into the mammary fat pad of female BALB/c mice resulted in robust tumor growth throughout the duration of the study, consistent with previous reports using the same mammary carcinoma cell line [34; 74]. No differences were observed between baseline measurements of mechanical paw withdrawal thresholds and those taken on the day tumors were first detected. Although inoculation of 4T1 cells can induce cancer-associated pain in mice independent of chemotherapy, our study did not detect such alterations within the experimental timeline. This discrepancy may reflect to the shorter observation window post-inoculation in our study [13].

Paclitaxel treatment induced both mechanical and cold hypersensitivity in mice bearing mammary tumors. Acute administration of AB120 alleviated paclitaxel-induced behavioral hypersensitivities throughout the study period, while the inactive isotype control, AB100, had no effect. Small-molecule CB2 agonists reduce pain behaviors in models of cancer and chemotherapy-induced neuropathy. AM1241 and JWH015 suppress cancer-related pain in murine models of bone cancer [12; 13; 41; 68]. Similarly, monoclonal antibodies targeting key mediators of pain processing, such as nerve growth factor (NGF) and vascular endothelial growth factor A (VEGF-A), attenuate chronic cancer pain and CIPN in preclinical models [4; 31; 46; 72]. Further studies are required to assess effects of extended treatment schedules with CB2-specific antibody agonists on paclitaxel-induced behavioral sensitivities and tumor growth dynamics in mice bearing mammary tumors.

Acute treatment with AB110 and AB120 did not induce motor ataxia or impair locomotion in the rotarod and activity meter tests, respectively. By contrast, the CB1/CB2 mixed agonist CP55-940 increased rest time and reduced total distance traveled. These findings are consistent with previous studies demonstrating that small-molecule CB2 agonists do not typically produce locomotor deficits [15; 56], whereas CB1 activation can lead to various unwanted side effects, including lethargy [14; 26]. In fact, CB2 agonists may enhance motor function in models of neurological dysfunction [52; 53]. These effects are consistent with limited expression of CB2 receptors in the brain, but participation of these inducible receptors in brain function remains unclear, especially given limitations on accurate detection of receptor expression [69]. Altogether, our findings highlight the favorable safety profile of CB2-specific antibody agonists for the treatment of CIPN.

*In vitro*, paclitaxel-induced cytotoxicity in mouse tumor cells (4T1) was preserved when in combination with CB2-specific antibody agonists in the MTT assay. In addition, potential additive effects were detected with the combination of paclitaxel and CB2 antibody agonists in non-tumor HEK-293 cell viability, even though each drug by itself did not produce significant HEK-293 cell toxicity. Small molecule CB2 agonists modulate breast cancer cell viability and metastasis potential [27]. CB2 agonists, such as MDA7, did not interfere with the cytotoxicity of paclitaxel in cell viability assays using different human breast cancer cell lines [48]. Therefore, CB2-specific antibody agonists show potential to treat CIPN, without altering the effectiveness of chemotherapy agent paclitaxel in producing tumor cell cytotoxicity *in vitro*. More work is necessary to extend these observations to tumor bearing mice receiving prophylactic dosing with therapeutic antibodies.

In conclusion, this study presents the first evidence of antinociceptive efficacy of first in class therapeutic antibody agonists targeting a GPCR—specifically CB2 receptors—in a mouse neuropathy model. CB2-specific antibody agonists, but not an isotype control, produce a sustained attenuation of paclitaxel-induced mechanical and cold hypersensitivity, which are hallmark sensory symptoms of CIPN. Our findings also underscore the safety and tolerability of these compounds with repeated administration, showing no signs of analgesic tolerance or locomotor impairment. Furthermore, efficacy of the lead antibody agonist AB120 was CB2-mediated and retained in 4T1-tumor bearing mice that were treated with paclitaxel to induce CIPN, enhancing translational relevance.

## Acknowledgements

This research was primarily supported by R43 CA241513 and R44 CA241513 (LS and AGH). CB2 KO mice were provided with support of DA047858 (to AGH and Ken Mackie). JLW and ES are supported by NIDA T32 training grant DA024628.

## Supplemental Figure Legends

**Supplemental Figure 1.**
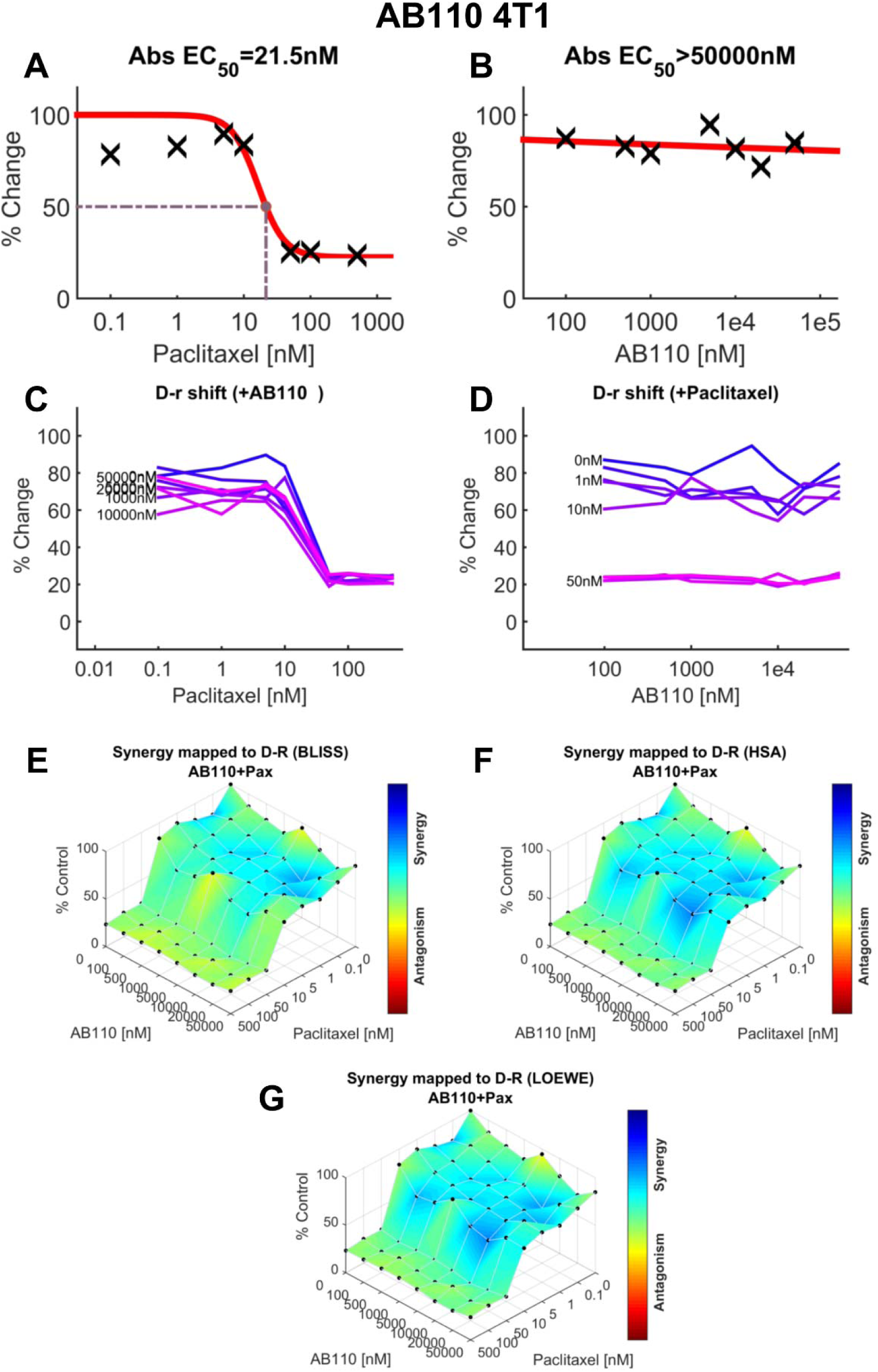
AB110 and paclitaxel produce additive effects in 4T1 cells to reduce murine mammary carcinoma cell line viability. (**A**) Dose-response for the effect of paclitaxel in 4T1 cell viability (EC50=21.5 nM). (**B**) Dose-response curve for the effect of AB110 in 4T1 cell viability (EC50>50 µM). (**C**) Paclitaxel dose response shift observed in the presence of increasing concentrations of AB110. (**D**) AB110 dose response shift observed in the presence of increasing concentrations of paclitaxel in 4T1 cell viability. (**E–G**) The 3-dimensional landscape of the dose matrix is represented on a color scale, where blue reflects evidence of synergy and red reflects evidence of antagonism. The landscape of the dose matrix of combination responses for AB110 and paclitaxel based on the (**E**) Bliss model, (**F**) Highest Single Agent (HSA) model and (**G**) Loewe model. Each model supports additive effects of the combination in reducing 4T1 tumor cell line viability. Cell viability is plotted as % control. (n = 3 experiments).

**Supplemental Figure 2.**
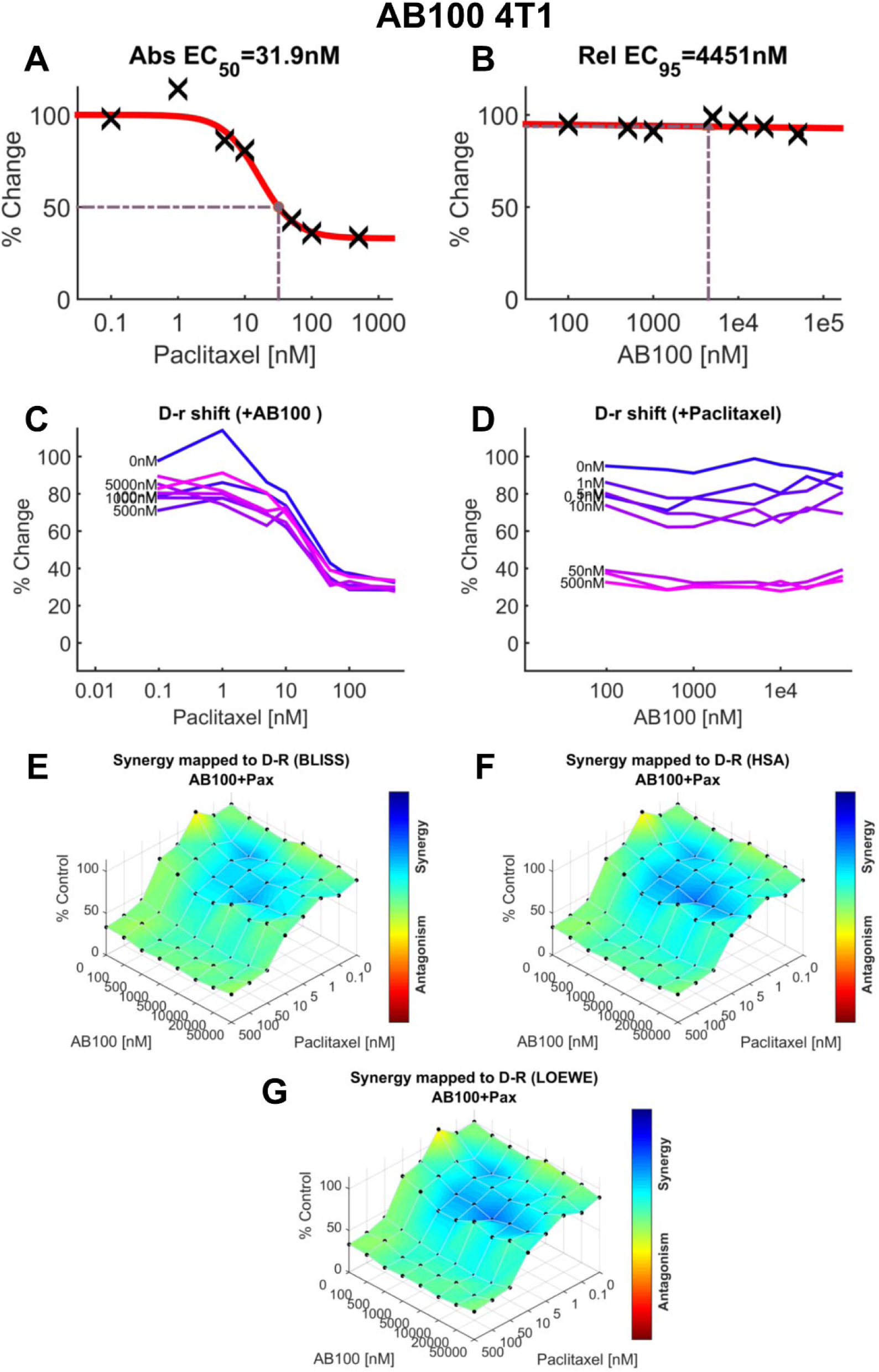
AB100 produces slight antagonism or additive effects with paclitaxel in 4T1 cells to reduce murine mammary carcinoma cell line viability. (**A**) Dose-response for the effect of paclitaxel on 4T1 cell viability (EC50=31.9 nM). (**B**) Dose-response curve for the effect of AB100 in 4T1 cell viability (EC95=4.451 µM). (**C**) Paclitaxel dose response shift observed in the presence of increasing concentrations of AB100 in 4T1 cell viability. (**D**) AB100 dose response shift observed in the presence of increasing concentrations of paclitaxel in tumor cell viability (4T1). (**E–G**) The 3-dimensional landscape of the dose matrix is represented on a color scale, where blue reflects evidence of synergy and red reflects evidence of antagonism. The landscape of the dose matrix of combination responses for AB100 and paclitaxel based on the (**E**) Bliss model, (**F**) Highest Single Agent (HSA) model and (**G**) Loewe model. Each model supports either modest antagonism or additive effect of the combination in reducing 4T1 tumor cell line viability. Cell viability is plotted as % control. (n = 3 experiments).

**Supplemental Figure 3.**
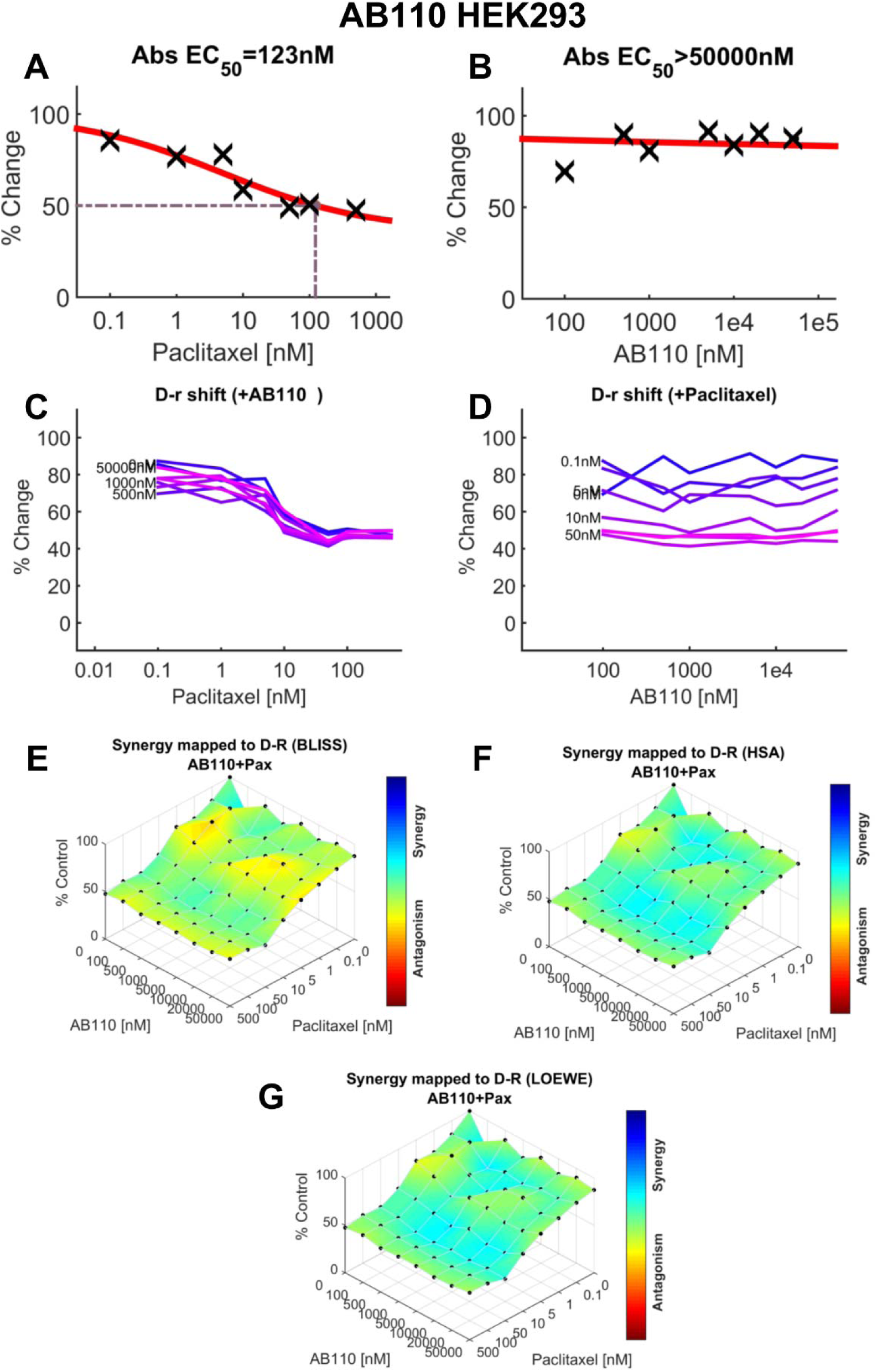
Additive effects of AB110 and paclitaxel on nontumor HEK293 cells. (**A**) Dose-response for the effect of paclitaxel on viability of HEK293 cells (EC50=123 nM). (**B**) Dose-response curve for the effect of AB110 on viability of HEK293 cells (EC50>50 µM). (**C**) Paclitaxel dose response shift observed in the presence of increasing concentrations of AB110 in non-tumor cell viability (HEK293). (**D**) AB100 dose response shift observed in the presence of increasing concentrations of paclitaxel in non-tumor cell viability. (**E–G**) The 3-dimensional landscape of the dose matrix is represented on a color scale, where blue reflects evidence of synergy and red reflects evidence of antagonism. The landscape of the dose matrix combination responses for AB110 and paclitaxel based on the (**E**) Bliss model, (**F**) Highest Single Agent (HSA) model and (**G**) Loewe model. Cell viability is plotted as % control (n = 4 experiments).

**Supplemental Figure 4.**
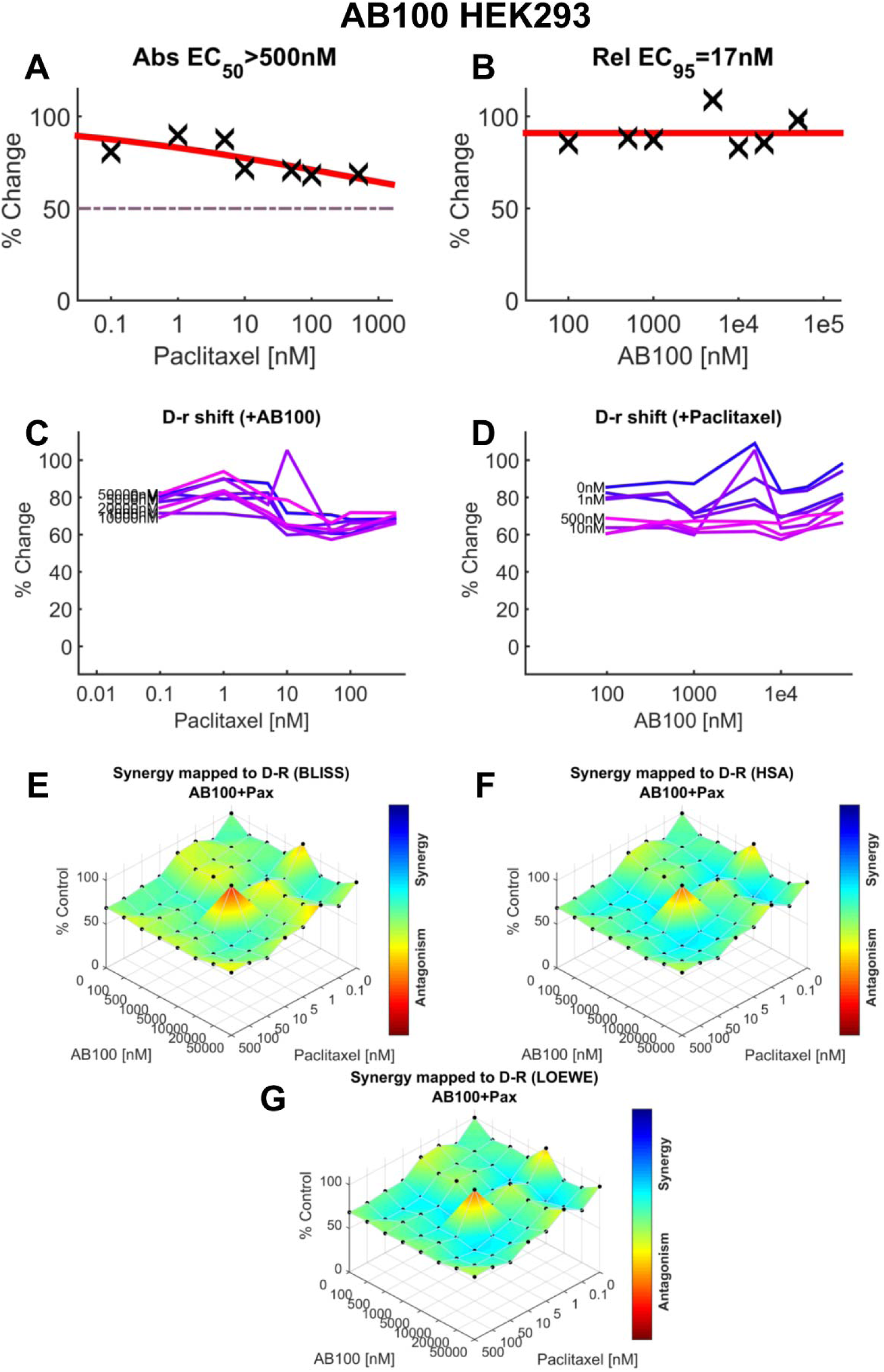
Additive effects of AB100 and paclitaxel on the viability of non-tumor HEK293 cells. (**A**) Dose-response curve for the effect of paclitaxel on viability of HEK293 cells (EC50>500 nM). (**B**) Dose-response for the effect of AB100 on viability of HEK293 cells (EC95=17 nM). (**C**) Paclitaxel dose response shift observed in the presence of increasing concentrations of AB100. (**D**) AB100 dose response shift observed in the presence of increasing concentrations of paclitaxel. (**E–G**) The 3-dimensional landscape of the dose matrix is represented on a color scale, where blue reflects evidence of synergy and red reflects evidence of antagonism. The landscape of the dose matrix combination responses for AB100 and paclitaxel based on the (**E**) Bliss model, (**F**) Highest Single Agent (HSA) model and (**G**) Loewe model. Cell viability is plotted as % control (n = 4 experiments).

**Supplemental Table 1.**
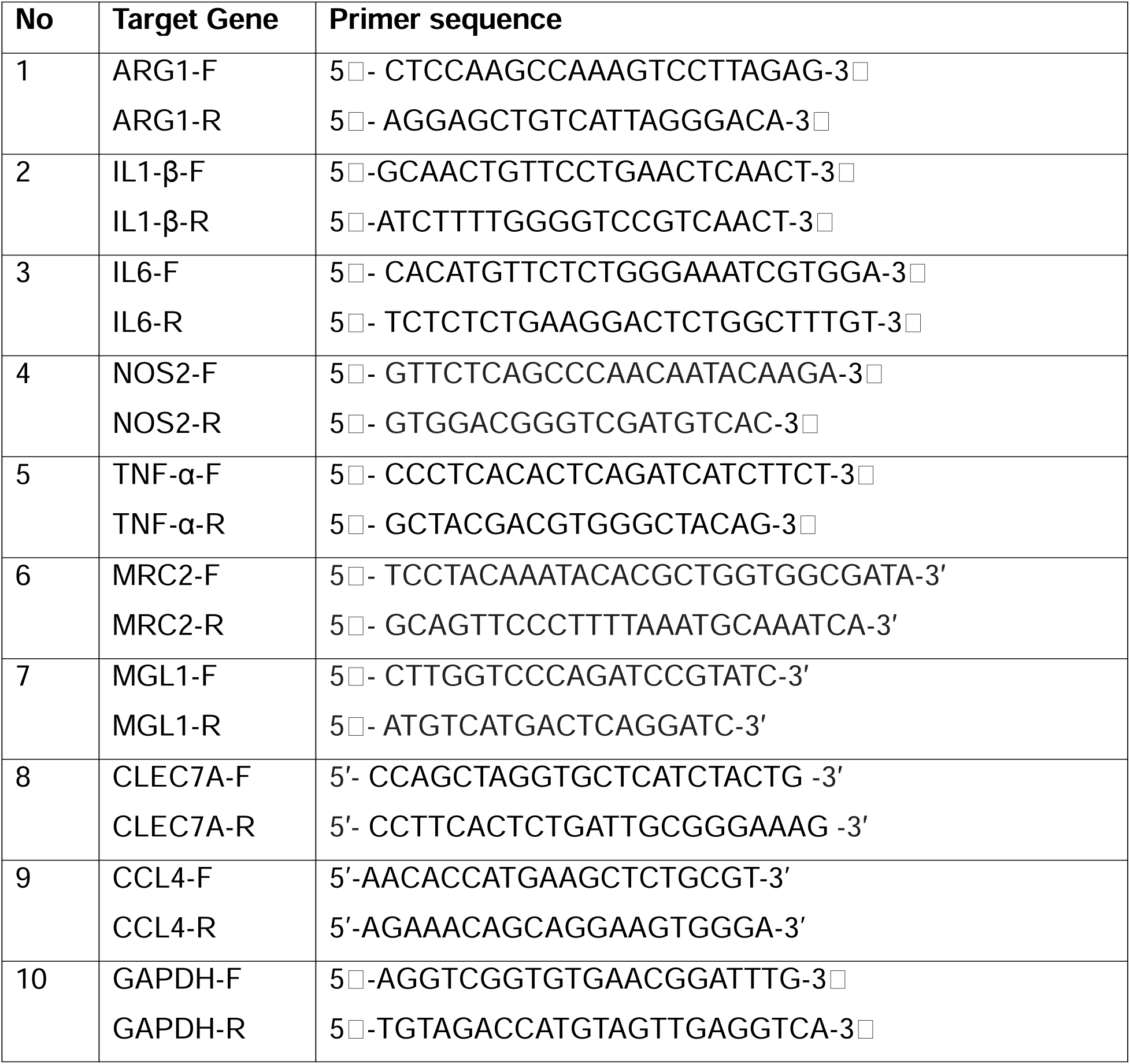
Primers used in real-time PCR.

**Supplemental Table 2.**
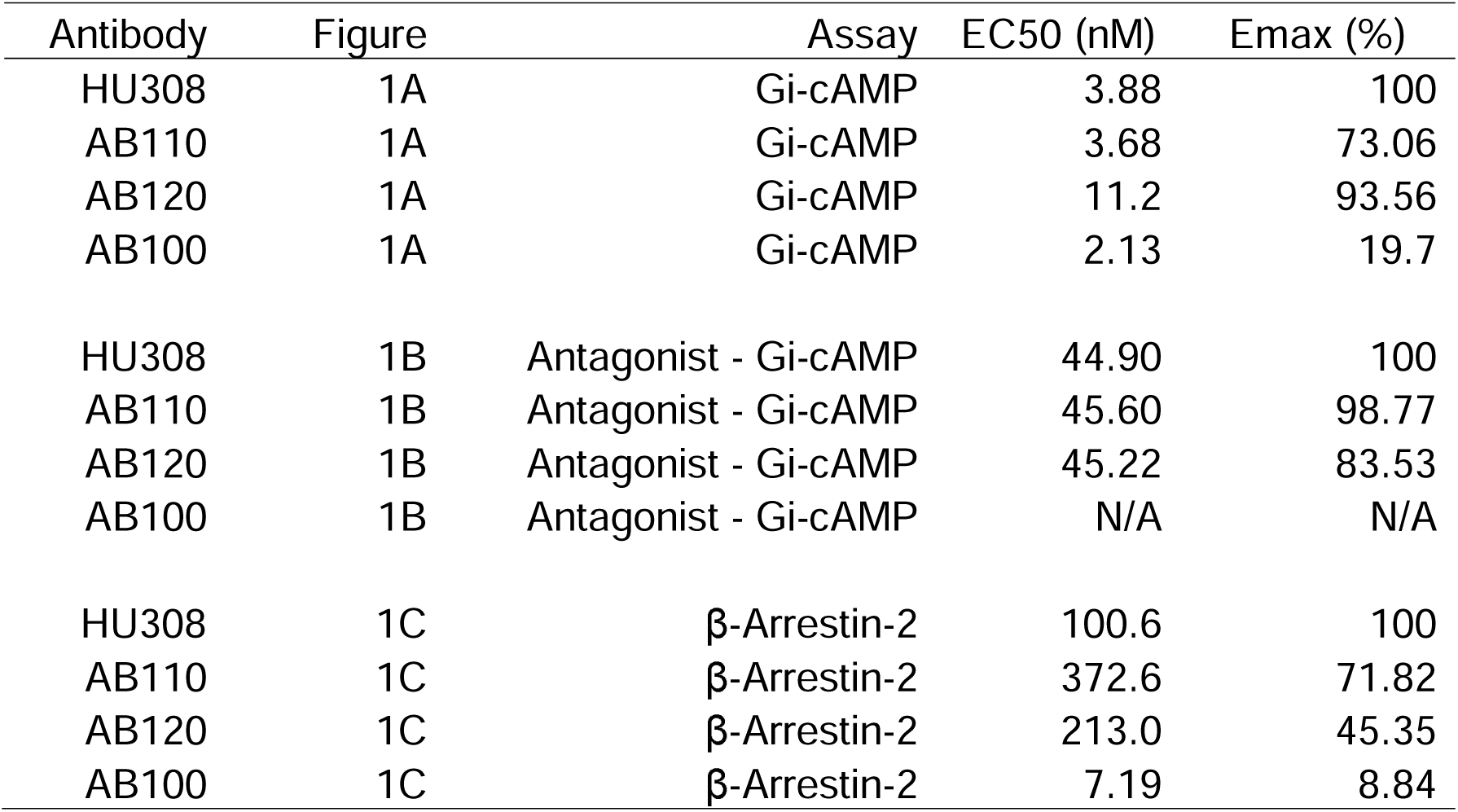
Efficacy and potency profiles of CB2 antibody agonists.

| Antibody | Figure | Assay | EC50 (nM) | Emax (%) |
| --- | --- | --- | --- | --- |
| HU308 | 1A | Gi-cAMP | 3.88 | 100 |
| AB110 | 1A | Gi-cAMP | 3.68 | 73.06 |
| AB120 | 1A | Gi-cAMP | 11.2 | 93.56 |
| AB100 | 1A | Gi-cAMP | 2.13 | 19.7 |
| HU308 | 1B | Antagonist - Gi-cAMP | 44.90 | 100 |
| AB110 | 1B | Antagonist - Gi-cAMP | 45.60 | 98.77 |
| AB120 | 1B | Antagonist - Gi-cAMP | 45.22 | 83.53 |
| AB100 | 1B | Antagonist - Gi-cAMP | N/A | N/A |
| HU308 | 1C | β-Arrestin-2 | 100.6 | 100 |
| AB110 | 1C | β-Arrestin-2 | 372.6 | 71.82 |
| AB120 | 1C | β-Arrestin-2 | 213.0 | 45.35 |
| AB100 | 1C | β-Arrestin-2 | 7.19 | 8.84 |

**Supplemental Table 3.**
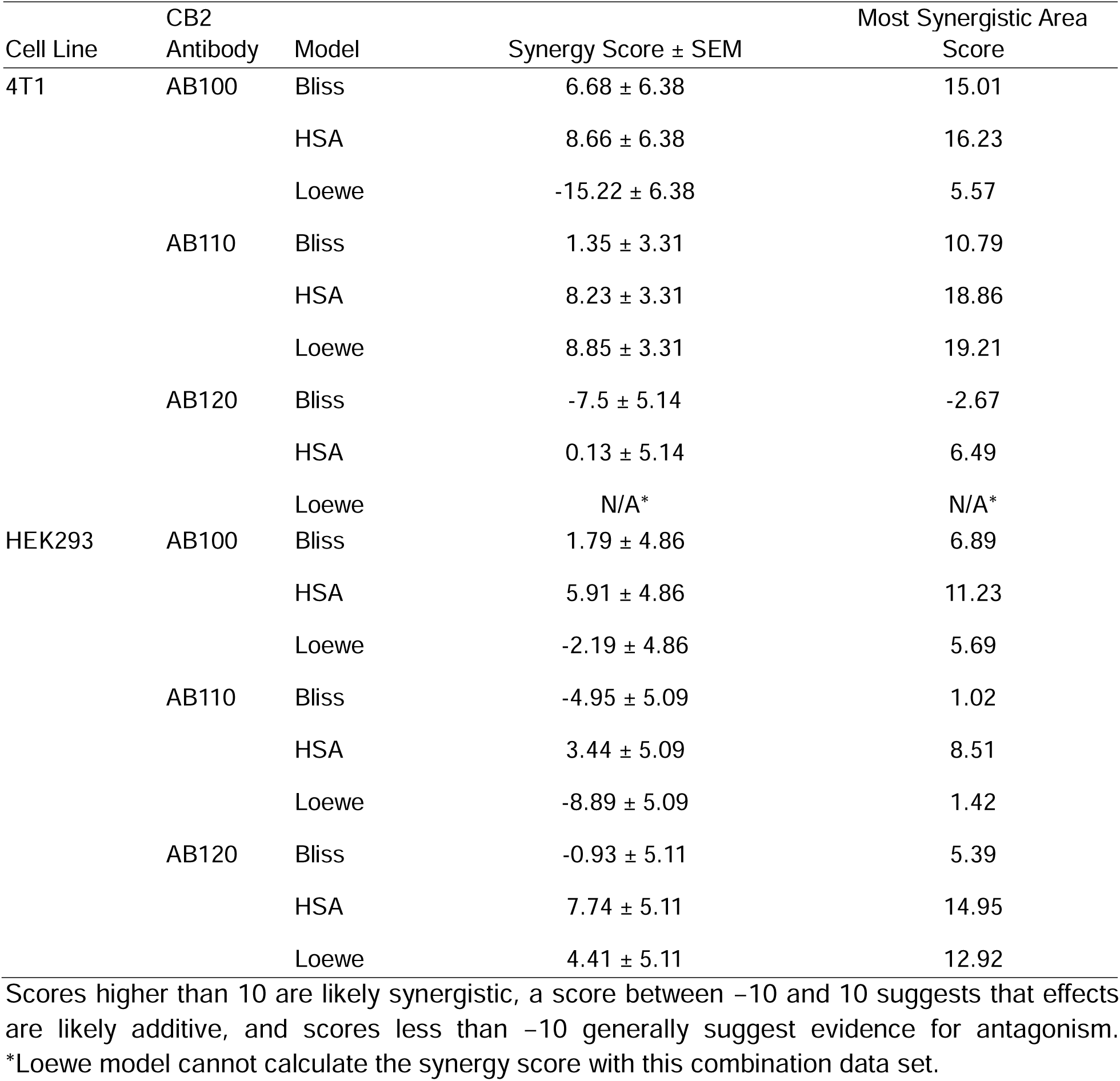
Synergy score to quantify degree of synergy for paclitaxel and CB2-specific antibody agonists.

